# Blocking somatic repeat expansion and lowering huntingtin via RNA interference synergize to prevent Huntington’s disease pathogenesis in mice

**DOI:** 10.1101/2025.06.24.661398

**Authors:** Jillian Belgrad, Ashley Summers, Christian Landles, Jonathan R. Greene, Samuel Hildebrand, Emily Knox, Ellen Sapp, Nozomi Yamada, Raymond Furgal, Rachael Miller, Georgina F. Osborne, Kathryn Chase, Eric Luu, Jason Freedman, Brianna Bramato, Nicholas McHugh, Vicky Benoit, Daniel O’Reilly, Paul Greer, Gillian P. Bates, Thomas F. Vogt, Ramee Lee, David Howland, Marian DiFiglia, Neil Aronin, Anastasia Khvorova

## Abstract

Huntington’s disease (HD) is a progressive neurodegenerative disorder with no approved therapies. Two major molecular drivers—somatic expansion of inherited CAG repeats and toxic mutant HTT (mHTT) variants—lead to neuronal dysfunction. Despite multiple trials, HTT-lowering strategies have not shown meaningful clinical benefit. Using therapeutic divalent siRNAs, we assessed the long-term impact of silencing MSH3 (a key regulator of somatic expansion), HTT, or both. In Q111 HD mice (>110 CAGs), which exhibit robust expansion, mHTT inclusions, and transcriptional dysregulation by 12 months, long-term MSH3 silencing blocked expansion, reduced inclusions, and reversed gene expression changes. HTT silencing alone had limited effect, but combined MSH3/HTT targeting synergistically eliminated inclusions and restored transcriptomic profiles. Parallel treatment in wild-type mice showed no toxicity, supporting the safety of long-term intervention. These findings position somatic expansion as a promising therapeutic target and demonstrate the potential of RNAi-based co-silencing of MSH3 and HTT as a disease-modifying strategy for HD.

## INTRODUCTION

Huntington’s disease (HD) is a fatal, autosomal dominant condition characterized by neurodegeneration, particularly in the striatum, leading to chorea, motor impairment, and cognitive decline that typically manifests between the ages of 30 and 50. Currently, there are no disease-modifying treatments for HD patients.

HD is caused by inheriting 40 or more CAG trinucleotide repeats in exon 1 of the huntingtin (*HTT*) gene, which encodes a polyglutamine-rich mutant HTT protein. The presence of neuronal inclusions and mHTT protein aggregates is a hallmark of HD^1–4^. In mouse models, increased count and size of focal aggregates correlate with more advanced disease^4^. The role of mHTT aggregates as a driver of disease remains debated^1, 3^. Nevertheless, reduction of total HTT (wild-type and mutant) has been the main therapeutic approach explored over the past decade^5, 6^. Clinical trials for HTT-lowering drugs have faced challenges^7^, and demonstrating efficacy remains a work in progress.

Growing evidence indicates that a more complex interplay of cellular processes mediates HD pathogenesis^8^. Inherited CAG repeat length in germline cells influences the age of clinical diagnosis^9,10^. However, CAG repeat length in somatic cells, particularly in transcriptionally active neurons, is unstable, leading to further expansion over time^11–13^. This process, termed somatic repeat expansion, has been observed in patients decades before disease onset^14^, and can result in 300 or more CAG repeats by the advanced stages of HD^15, 16^. Somatic repeat expansion occurs in a mosaic manner, with striatal medium spiny neurons being most impacted^15, 16^. This expansion is associated with increased production of HTT1a^15^, a toxic product of aberrant *HTT* mRNA processing that uses early cryptic poly-adenylation sites and seeds mHTT aggregates found in human patients and mouse models^17, 18^. Somatic expansion is facilitated by the mismatch repair (MMR) pathway^12, 19, 20^. Silencing key MMR genes, particularly knockout or knockdown of MutS homolog 3 (*MSH3*), MutL homolog 1 (*MLH1*), MutL homolog 3 (*MLH3*), and post meiotic segregation increased 1 (*PMS1*), can halt somatic expansion in models of HD^12, 20–24^. With a better understanding of interacting pathways in HD—mHTT aggregation, somatic expansion, and HTT1a formation—efforts to identify disease-modifying therapies are ongoing.

An emerging threshold model of HD suggests that while inheriting :240 CAG repeats is enough to predict disease onset, mHTT may not be inherently toxic in the central nervous system (CNS) until CAG repeats exceed 150^8, 15, 22, 25–27^; however, the exact threshold in humans remains unclear. This threshold model proposes that blocking somatic expansion could be disease-modifying if CAG lengths are stabilized between 40 and 150, identifying a therapeutic window _for HD8,15,22,25,26._

Here, we used a therapeutically relevant modality of chemically stabilized divalent small interfering RNA (siRNA) to test whether blocking somatic expansion alone or in combination with HTT lowering can modify HD^28^. siRNAs are an established class of nucleic acid-based therapeutics, with seven siRNA drugs approved for liver and systemic use by the US Food and Drug Administration (FDA)^29, 30^. Divalent siRNA enable potent (>80%), sustained (at least 4-6 months), and non-toxic gene silencing in the central nervous system (CNS)^28^. Divalent siRNA-mediated silencing of *HTT* and *Msh3* have been preclinically validated^23, 28^, with divalent siRNA-mediated *Msh3* silencing shown to block somatic expansion *in vivo*^23^. We injected 2-month-old *Hdh*^Q111^ mice (which possess an individual germline range from 110- to 120-CAG repeat tracts) with chemically stabilized divalent siRNAs targeting *Msh3*, *HTT*, or both, and assessed somatic instability, HTT protein species expression and aggregation, neurodegenerative transcriptomic signatures, and behavior by 12 months of age.

Blocking somatic expansion alone prevented the formation of mHTT inclusions, improved nesting behavior, and reversed a significant portion of the neurodegenerative transcriptomic signature. HTT lowering (pan-allelic and without engaging HTT1a) had a minimal impact on these outcomes. However, blocking somatic expansion in combination with HTT lowering completely abolished mHTT inclusions, reduced levels of aggregated HTT1a, markedly improved nesting behavior, and synergistically reversed the neurodegenerative HD transcriptomic signature more effectively than either strategy alone. These findings demonstrate that blocking expansion with therapeutic agents can slow HD pathogenesis, even in vulnerable cells that have achieved >110 CAGs. Our results also suggest that HTT lowering can be modestly disease-modifying when combined with approaches that block somatic expansion. Combination therapy in cases with >120 CAGs may be more effective than either approach alone. This work informs a potential disease-modifying therapeutic strategy for HD.

## RESULTS

### Divalent siRNA targeting *Msh3*, with or without *HTT* co-silencing, suppressed somatic expansion, whereas *HTT* silencing alone reduced *HTT* levels without affecting expansion in 12-month-old *HdhQ111* mice

To explore whether blocking somatic repeat expansion can slow or prevent HD pathogenesis, we designed a long-term study in wild-type (WT) mice and the *Hdh*^Q111^ mouse model (hereafter referred to as Q111 mice), outlined in Figure 1a. Q111 mice are a knock-in model with mutant human *HTT* exon 1 replacing mouse exon 1 in the mouse *Htt* gene. From 2 month of age, Q111 mice undergo measurable somatic repeat expansion and demonstrate robust neurodegenerative signatures in the striatum and cortex, whilst at 6 months of age, Q111 mice exhibit mHTT aggregation in the striatum^31–35^. WT and Q111 mice were treated with artificial cerebrospinal fluid (aCSF)^36^, divalent siRNA programmed with a non-targeting control (NTC) sequence or sequences targeting *Msh3* (10 nmol/10µL, 240 ng) or *HTT* (10 nmol/10µL, 240 ng), or a mixture of divalent siRNA targeting *Msh3* and divalent siRNA targeting *HTT* (10 nmol each) via intracerebroventricular (ICV) injection at 2 months of age. For all cohorts, mice were reinjected with their respective treatment at 5 and 8 months of age and sacrificed at 12 months of age. The divalent siRNA scaffold and chemical modification are shown in Fig. 1b,c. The divalent siRNAs (sequences shown in Supplemental Table 1) were previously validated and cross-react with both human and mouse mRNA^23, 28^. The HTT sequence used was pan-allelic for WT and mutant *HTT*^28^ and targeted the full-length *HTT* 3’UTR therefore does not engage HTT1a.

**Fig. 1:**
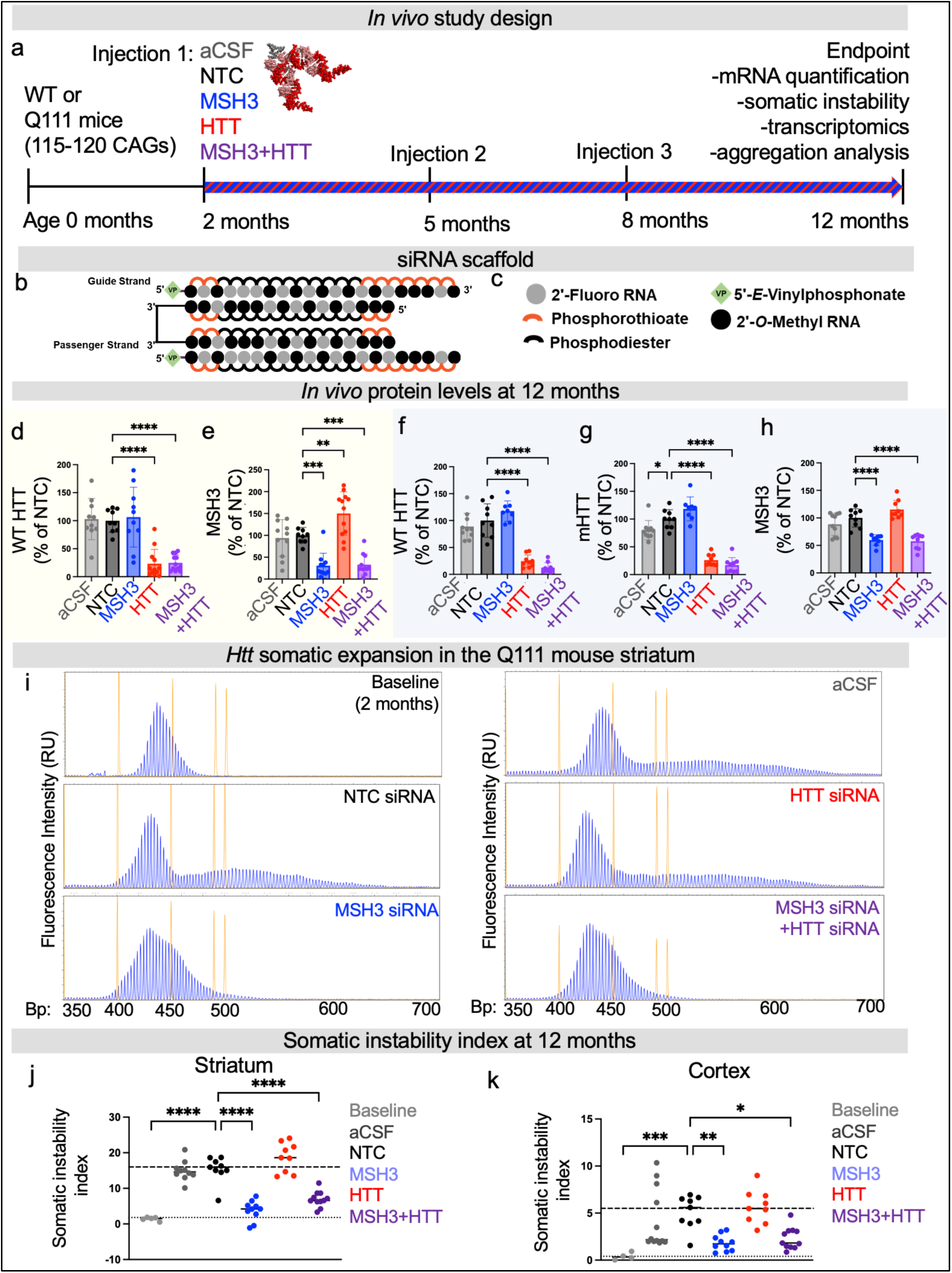
Divalent siRNA silence target gene expression and block somatic repeat expansion in Q111 mice. (**a**) In vivo study design and treatment paradigm. 2-month-old wild-type (WT) or Q111 mice were treated with artificial CSF (aCSF), or divalent siRNA programmed with sequences targeting a non-targeting control (NTC), *MSH3, HTT,* or *MSH3* and *HTT*, and euthanized at 12 months old. (**b**) Divalent siRNA scaffold and (**c**) chemical modification pattern key. (**d, e)** Endpoint protein levels in WT striatum: (**d**) WT HTT, (**e**) MSH3. (**f-h**) Endpoint protein levels in Q111 striatum: (**f**) WT HTT, (**g**) mutant HTT, (**h**) MSH3. (**i**) Representative striatum fragment analysis curves for baseline (2 months Q111 untreated) or Q111 mice (12 months old) treated with artifical CSF (aCSF), non-targeting control (NTC) siRNA, HTT siRNA, MSH3 siRNA, or MSH3+HTT siRNA combination. (**j,k**) Somatic instability index quantified from fragment analysis curves in (**j**) striatum or (**k**) medial cortex. WT cohorts: aCSF, n=12; NTC, n=10; MSH3 siRNA, n=12; HTT siRNA, n=12; MSH3+HTT siRNA, n=12 mice. Q111 cohorts: aCSF, n=12; NTC, n=10; MSH3 siRNA, n=10; HTT siRNA, n=9; MSH3+HTT, n=11 mice. Statistics are one-way ANOVA with Dunnett’s multiple comparisons test. * p < 0.05; ** p < 0.01; *** p < 0.001; **** p < 0.0001.

Following euthanasia, target mRNA and protein knockdown were assessed in harvested brain regions using branched DNA (Quantigene) assay or western blot, respectively. Target expression was robustly silenced at the experimental endpoint with divalent siRNA achieving 40-70% *HTT* and 20-50% *Msh3* mRNA lowering across the intended WT and Q111 treatment cohorts (Supplementary Fig. 1). At the protein level, divalent siRNA achieved 70-90% HTT and 50-70% MSH3 silencing in the intended WT (Fig. 1d,e) and Q111 (Fig. 1f-h) treatment cohorts (Fig. 1d-h, Supplementary Fig. 2).

We next assessed the somatic expansion of the CAG trinucleotide repeat within the *HTT* locus using fragment analysis. Fragment analysis curves were quantified using a somatic instability index, which determines the skewness of CAG lengths relative to the main CAG length^37^ (Fig. 1i-k). Representative fragment analysis curves of 12-month-old Q111 mice with their respective siRNA treatments are shown in Fig. 1i. Baseline CAG lengths were measured in the striatum of 2-month-old Q111 mice, revealing a somatic instability index of 1.4 ± 0.45. By 12 months of age, aCSF- and NTC-treated Q111 mice had striatal somatic instability indices of 14.9 ± 2.7 and 15.4 ± 3.4, respectively (Fig. 1j, baseline vs NTC, p < 0.0001), indicating significant somatic expansion. At 12 months, the striatal somatic instability indices of Q111 mice treated with divalent siRNAs targeting *Msh3* alone or with *HTT* were 3.7 ± 2.6 and 6.9 ± 2.1, respectively (NTC vs MSH3, p < 0.0001; NTC vs MSH3+HTT, p < 0.0001), indicating expansion was blocked following MSH3 knockdown. This finding is consistent with prior work^23^. siRNA-mediated lowering of HTT had no impact on somatic expansion (somatic instability index of 18.7 ± 3.9; NTC vs HTT, p = 0.086), which is also consistent with prior work^23, 38^. Similar results were observed in the medial cortex (Fig. 1k, NTC vs. MSH3, p < 0.01; NTC vs. MSH3+HTT, p < 0.05), a brain region with a lower degree of overall somatic expansion.

### Blocking expansion reduces focal aggregates of mHTT and is enhanced with HTT co-lowering

With somatic expansion blocked and expanded HTT reduced in the target cohorts, we next examined how therapeutic modulation of MSH3 or HTT affected HD outcomes. To first assess mHTT load we performed immunohistochemistry in the same cohort of mice using the monoclonal EM48 antibody (Fig. 2)^3^, which binds an epitope that has been mapped within the last 12 amino acids of HTT1a and detects aggregated mHTT^39^. Previous studies have shown that EM48+ aggregates start diffusely and become more focal (i.e. inclusions) as the disease progresses^40^. To evaluate disease progression in treated Q111 mice, we quantified inclusions, diffuse staining, and the ratio of inclusion-to-total or diffuse-to-total EM48+ events.

**Fig. 2.**
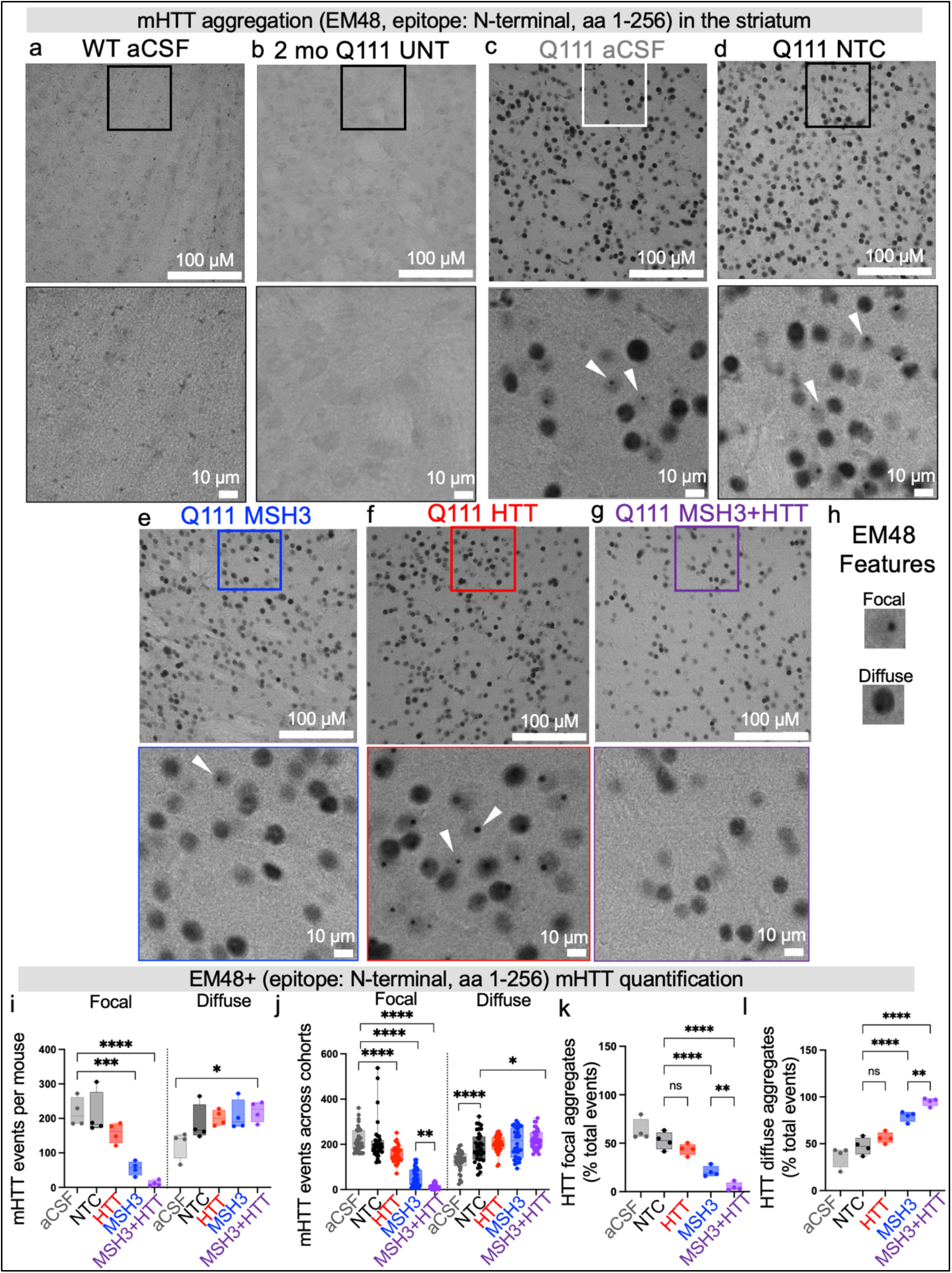
Blocking expansion eliminates focal EM48+ mHTT puncta formation, enhanced mHTT puncta lowering with HTT co-lowering. (**a-g**) Striatal sections probed with EM48 immunohistochemistry. Representative images shown. The top panel is 40x, and the bottom panel is an area within a boxed region of interest. White arrow indicates representative EM48+ focal puncta. (**a**) WT aCSF (**b**) 2-month-old untreated Q111 (**c**) aCSF treated Q111 (**d**) NTC treated Q111 (**e**) MSH3 siRNA treated Q111 (**f**) HTT siRNA treated Q111 (**g**) MSH3+HTT combination siRNA treated Q111 (**h**) Representative histological characterization of puncta (top) or diffuse (bottom) EM48+ mHTT events (**i**) Quantified puncta (right) or diffuse (right) events per mouse totaled from three 40x images across three 40 µM sections (nine images counted per mouse). Each dot is average from one mouse. (**j**) mHTT puncta (left) or diffuse (right) quantified over each cohort. Same data as panel i, but each dot is puncta or diffuse EM48+ event from one 40x image, not averaged per mouse. (**k**) Percentage of focal puncta to total EM48+ events. Each dot is the average of nine slides from one mouse. (**l**) Percentage of diffuse EM48+ events to total EM48+ events. Each dot is the average of nine slides from one mouse. Total EM48+ events are defined as focal plus diffuse events. n=4 mice/group strained and quantified. Statistics are one-way ANOVA with Holm-Šídák’s multiple comparisons test. * p < 0.05; ** p < 0.01; *** p < 0.001; **** p < 0.0001.

No EM48+ reactivity was detected in WT mouse striatum (Fig. 2a). In 2-month-old Q111 mice, we observed slight diffuse EM48+ staining (Fig. 2b)^4^. In the 12-month-old Q111 mice treated with aCSF or NTC siRNA, we observed robust EM48+ stains, diffuse and inclusions (Fig. 2c, d, h,i). aCSF- and NTC-treated Q111 mice have an average inclusion count of 217 ± 36 and 210 ± 55 per field, respectively. Q111 mice treated with divalent siRNA targeting *HTT* had an average inclusion count of 158 ± 25, which was not significantly different from NTC (Fig. 2f, i, NTC vs. HTT, p = 0.1884), suggesting that silencing HTT alone had no significant impact on mHTT aggregation. Q111 mice treated with divalent siRNA targeting *Msh3* had a significantly reduced average number of inclusions, 55 ± 7, compared to NTC control (Fig. 2e, i, NTC vs. MSH3, p< 0.01). This reduction was even more pronounced in mice treated with divalent siRNA targeting both *Msh3* and *HTT*, which had an average inclusion count of 11 ± 7 (Fig. 2g, i, NTC vs. MSH3+HTT, p < 0.001). The enhancement observed with combined MSH3 and HTT treatment, compared to MSH3 alone, was especially pronounced when we analyzed the total counts in each cohort (Figure 2g,i,j; MSH3 vs MSH3+HTT, *p* = 0.0049).

In comparison, the diffuse EM48+ staining remained largely unchanged following siRNA treatment (Fig. 2h,i). The only notable difference was a slight but statistically significant increase in the number of diffuse aggregates in the HTT+MSH3 condition, which showed a significantly higher count compared to the aCSF control (Fig. 2i; NTC vs. MSH3+HTT, *p* < 0.001).

Together, these findings show that blocking somatic expansion, but not lowering HTT alone, selectively eliminates mHTT inclusions while preserving diffuse staining. This effect was evident both in per-mouse analyses and when compiling counts across cohorts. The inclusion count was even further reduced with siRNA combination treatment compared to either treatment alone (Fig. 2i,j).

To characterize mHTT and pathology progression, we determined the ratio of inclusions or diffuse staining (Fig. 2h) to the total EM48+ events (Fig. 2k,l). A decrease in the total number of EM48+ events were evident in Q111 cohorts treated with MSH3 or MSH3+HTT siRNA (Supplementary Fig. 3). In Q111 mice treated with aCSF or NTC, most EM48+ staining forms an inclusion (63 ± 9% and 53 ± 8%, respectively). In Q111 mice treated with divalent siRNA targeting *HTT*, inclusions make up 43 ± 5% of the EM48+ staining, which was not significantly different from controls (NTC vs. HTT, p = 0.076). In Q111 mice treated with divalent siRNA targeting *Msh3* with or without *HTT*, the prevalence of inclusions events significantly drops to 21 ± 5% or 5 ± 4, respectively (NTC vs. MSH3, p < 0.001; NTC vs. MSH3+HTT, p < 0.0001). While MSH3 treatment significantly reduced the pervalence of mHTT inclusions, combination treatment led to a significantly smaller percentage of mHTT inclusions than MSH3 alone (Fig 2k, MSH3 vs MSH3+HTT, p< 0.01). The inverse was true for the ratio of diffuse staining to total events. aCSF- and NTC-treated Q111 mice had few diffuse stains (36 ± 9% and 47 ± 8%, respectively). Treatment with divalent siRNA targeting HTT did not significantly change the ratio of diffuse staining in Q111 mice (56 ± 5%, NTC vs. HTT, p = 0.52). In contrast, treatment with divalent siRNA targeting *Msh3* resulted in a significantly higher ratio of diffuse staining in Q111 mice (79 ± 5%, NTC vs. MSH3, p <0.0001). The ratio of diffuse staining in Q111 mice treated with divalent siRNA targeting both *Msh3* and *HTT* was 94 ± 4%, a significantly higher ratio than those of mice treated with NTC or divalent siRNA targeting *Msh3* (NTC vs. MSH3+HTT, p <0.0001; MSH3 vs. MSH3+HTT, p<0.05), indicating significantly slower disease progression.

Overall, these results suggest that mHTT inclusions, but not diffuse mHTT, are sensitive to blocking somatic expansion. Inclusions are not sensitive to total, full length mutant HTT levels. Blocking expansion with interventional siRNA can significantly reduce mHTT inclusions, without impacting diffuse staining. mHTT inclusions were further reduced when blocking expansion was combined with HTT co-lowering indicating a shift toward a slower aggregation phenotype. Previous studies have shown that reducing HTT1a expression can eliminate diffuse mHTT^41^. Future work should explore whether a combination of blocking somatic expansion and lowering both HTT and HTT1a can fully eliminate pathological EM48+ reactivity in Q111 mice.

To further analyze the impact of blocking expansion on mHTT aggregates, we performed immunohistochemistry using the PHP1 antibody in control and divalent siRNA-treated Q111 mouse striatum. PHP1 binds to the proline-rich region at the end of the CAG repeat expansions. *Msh3* knockout in mice has been shown to eliminate PHP1+ inclusions in the striatum^22^. We also co-stained with DAPI and DARPP-32 to assess toxicity and localization, particularly in striatal medium spiny neurons and other nuclear cell populations within the striatum. We analyzed three key measures: (1) the number of inclusions, (2) the total area covered by aggregates in a field, and (3) the average size of each aggregate.

WT mice showed no reactivity with PHP1 and exhibited no detectable inclusions (Fig. 3a). In contrast, aCSF-treated and NTC-treated Q111 mice displayed a robust number of inclusions, averaging 337 ± 46 and 328 ± 57 per field, respectively (Fig. 3b,c,g). Treatment with divalent siRNA targeting *HTT* led to a similar number of inclusions, 262 ± 21 (Fig. 3d,g, NTC vs. HTT, p = 0.2166). Treatment with *Msh3-*targeting divalent siRNA resulted in a significant reduction to 34 ± 20 inclusions/field (Fig. 3e,g, NTC vs. MSH3, p < 0.0001). Similarly, silencing both MSH3 and HTT resulted in 16 ± 14 inclusions/field, a significant reduction compared to NTC control (Fig. 3f,g, p < 0.0001).

**Fig. 3.**
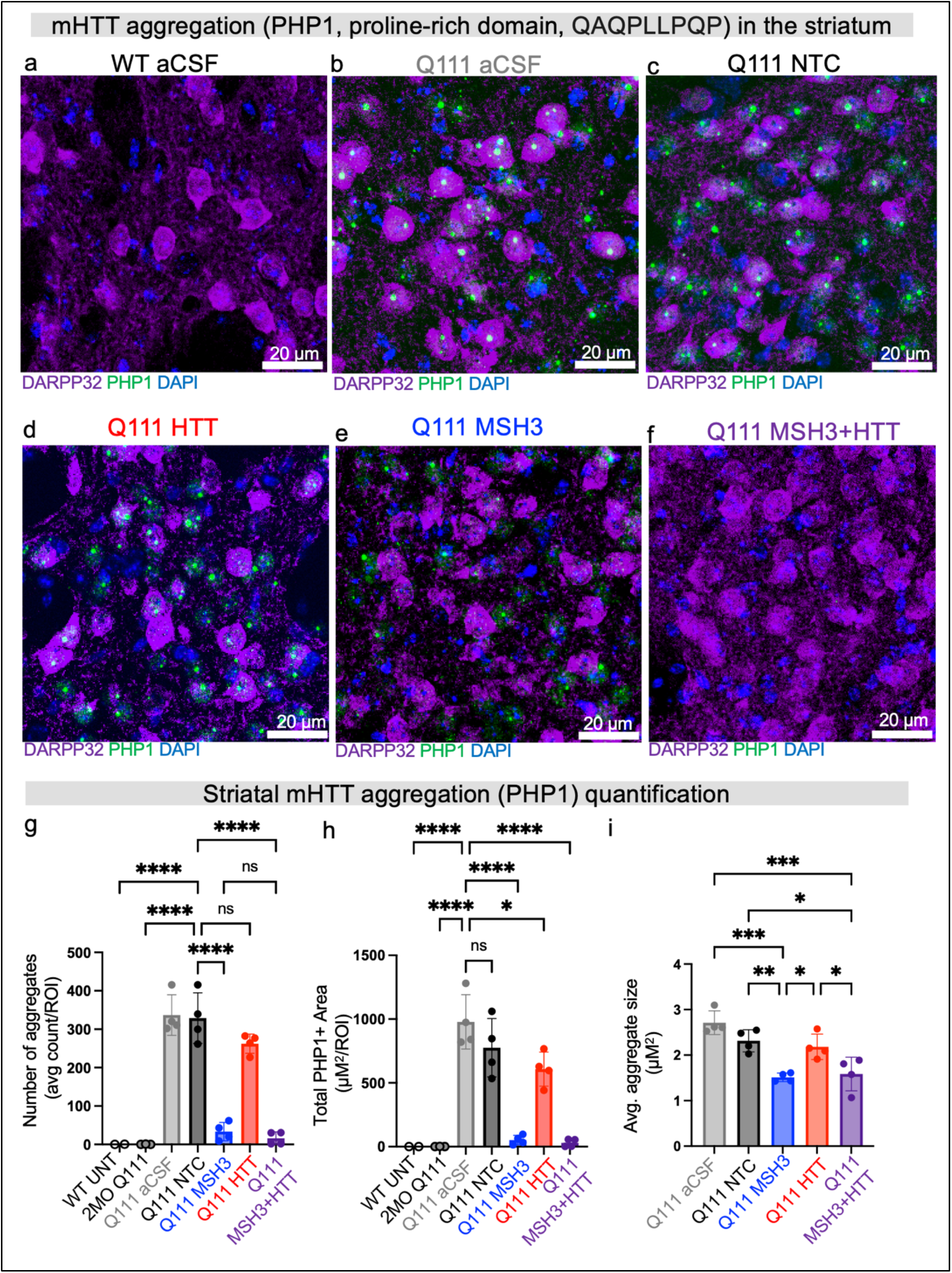
Blocking expansion eliminates PHP1+ mHTT aggregate formation in the Q111 mouse striatum. (a-f) Striatal immunofluorescence of PHP1 (green) co-stained with DARPP32 (purple) and DAPI (blue). Representative images of 12-month-old (**a**) aCSF-treated WT (**b**) aCSF treated Q111 (**c**) NTC treated Q111 (**d)** MSH3 siRNA treated Q111 (**e**) HTT siRNA treated Q111 (**f**) MSH3+HTT combination siRNA treated Q111 mice. Quantification of (**g**) average aggregate count per field of view, (**h**) PHP1+ area per ROI, (**i**) average aggregate size. n=4 mice/group imaged and quantified. Each dot summarizes events over six regions of interest, covering three striatal slices in one mouse. Statistics are one-way ANOVA with Šídák’s multiple comparisons test. * p < 0.05; ** p < 0.01; *** p < 0.001; **** p < 0.0001.

When analyzing the total area covered by inclusions, we observed 978 ± 184 µm² and 775 ± 198 µm² covered per field in aCSF- and NTC-treated mice, respectively (Fig. 3h). Whereas HTT lowering led to a slight reduction in the total area covered, measuring 608 ± 115 µm² (Fig. 3h, p = 0.0210 compared to aCSF; p = 0.0620 compared to NTC), lowering Msh3 alone or with HTT significantly reduced the area covered by inclusons, with 52 ± 32 µm² and 29 ± 26 µm² covered per field, respectively (Fig. 3h, p < 0.0001 for both comparisons to NTC).

The average size of inclusions in aCSF- and NTC-treated mice was 2.6 ± 2.6 µm² and 2.2 ± 2.02 µm², respectively. Silencing HTT led to a similar inclusion size (Fig. 3i, 2.5 ± 2.09 µm², p = 0.1315 vs. NTC). However, silencing MSH3 alone or with HTT led to inclusion sizes of 1.57 ± 1.43 µm² and 1.57 ± 1.89 µm², respectively, a significant reduction compared to NTC control (Fig. 3i, p = 0.0026 for NTC vs. MSH3; p = 0.0057 for NTC vs. MSH3+HTT). Collectively, these findings demonstrate a significant reduction in mHTT inclusion count and size following blockage of somatic expansion with or without co-reduction of HTT.

### Blocking somatic expansion increases soluble mHTT and, when combined with HTT lowering, reduces HTT1a aggregation

In the HD brain, HTT protein exists in different isoforms (e.g., WT, mutant, HTT1a) that can be soluble or aggregated^42^. The expression and aggregation of these species are associated with CAG length^2, 4, 17, 43, 44^. To determine the impact of blocking somatic expansion with or without co-reduction of HTT on HTT species expression and aggregation, homogenous time-resolved fluorescence (HTRF)^45, 46^ was performed on a separate cohort of untreated Q111 mice or Q111 mice treated with divalent siRNAs targeting an NTC, *MSH3, HTT*, or both *MSH3* and *HTT* (Fig. 4a, Supplemental Fig. 4a, Supplementary Table 1). Target knockdown and blocked expansion were validated in the expected treatment groups (Supplementary Fig. 4b-e). Striatal HTT protein species at 11 months are reported in Fig. 4, with results from all remaining brain regions and time points shown in Supplementary Fig. 5a-f. As expected, total full-length HTT (Fig. 4b, Supplementary Fig. 5a), endogenous mouse HTT (Supplementary Fig. 5b), and total full-length mutant HTT (Fig. 4c, Supplementary Fig. 5c) expression was reduced across brain regions in mice treated with siRNA targeting *HTT* alone or with dual *HTT* and *Msh3* (endogenous: NTC vs. HTT, p < 0.0001; NTC vs. MSH3+HTT, p< 0.000;1 full-length: NTC vs. HTT, p < 0.0001; NTC vs. MSH3+HTT, p< 0.0001).

**Fig. 4.**
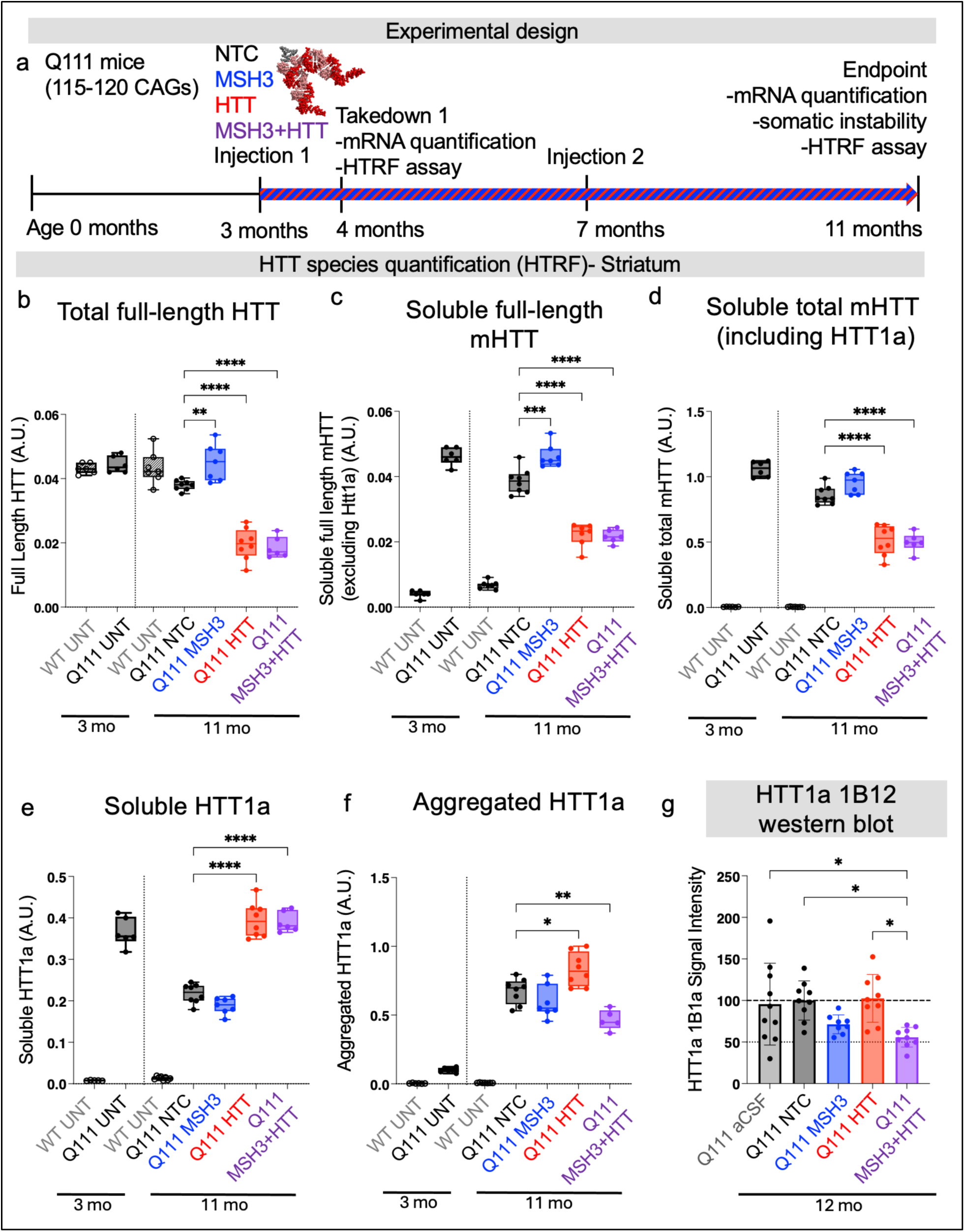
Blocking expansion maintains soluble mHTT levels; blocking expansion combined with lowered huntingtin reduces HTT1a aggregation. **(a)** Experimental design and treatments. (**b-f**) Striatal HTT protein species quantified using homogenous time-resolved fluorescence detected from respective antibody donor (D) or acceptor (A) pairs across time points and treatment groups. (**b**) Full-length total HTT (d: MAB2166-Tb, a: CHDI-1414-d2) (**c)** Soluble full-length mHTT (excludes HTT1a) (d: MAB2166-Tb, a: 4C9-488) (**d**) Total soluble mHTT (d: 2B7-Tb, a: MW1-d2) (**e**) Soluble HTT1a (d: 2B7-Tb, a: 11G2-d2) (**f**) Aggregated HTT1a (d:4C9-Tb, a:11G2-d2). Statistics are one-way ANOVA with Dunnett’s multiple comparisons versus NTC. Mice per group at two-month timepoint: WT n=6, Q111 n=6; Mice per group at 11-month endpoint: untreated WT n=7, NTC siRNA Q111 n=8, MSH3 siRNA Q111 n=7, HTT siRNA Q111 n=7, MSH3+HTT siRNA Q111 n=6. (**g**) Quantified western blot probing for HTT1a using antibody 1B12 in the striatum of the cohort of Q111 mice presented in Figure 1A. Statistics are one-way ANOVA with all comparisons and Tukey’s multiple comparison test. * p < 0.05; ** p < 0.01; *** p < 0.001; **** p < 0.0001.

Striatal soluble full-length mHTT levels declined over time in untreated or NTC-treated Q111 mice (Supplementary Fig. 5d, Q111 control at 2-, 4-, 7- vs 11-months), which is consistent with previous studies^4, 46^. At 11 months, Q111 mice treated with siRNA targeting *Msh3* had significantly higher soluble full-length mHTT (excludes HTT1a) (Fig. 4c, Supplementary 5c), suggesting shorter polyglutamine tracts (as a result of blocked somatic expansion) might decrease the propensity of mHTT for aggregation (Fig. 4c, NTC vs. MSH3, p < 0.001) and may represent a less advanced disease state. Although not statistically significant, full-length mHTT levels were also trending higher at the 4-month time point in these mice (Supplementary Fig. 5c).

The observed effects of blocking somatic expansion disappear when total mHTT (including HTT1a) is considered (Fig. 4d, Supplementary 5d, NTC vs. MSH3 p = 0.1626). This suggests that factors beyond the somatic expansion of CAG repeats at the *HTT* locus influence HTT1a levels as measured by HTRF. Divalent siRNA-mediated lowering of HTT alone or with Msh3 did lead to reduced total soluble mHTT (Fig. 4d, NTC vs. HTT, p< 0.0001; NTC vs. MSH3+HTT, p < 0.0001).

Soluble HTT1a is detectable as early as 2 months and declines with disease progression (Fig. 4e Supplementary Fig. 5e). By 11 months, soluble HTT1a was lower in Q111 mice treated with NTC and siRNA targeting *Msh3* compared to 2-month-old Q111 mice (Fig. 4e) because of recruitment into HTT aggregates. Treatment with Msh3-targeting siRNA led to a slight but nonsignificant decrease in soluble HTT1a compared to NTC control (p= 0.1626). HTT-lowering siRNA treatments maintained soluble HTT1a at high levels, suggesting that soluble HTT1a naturally decreases with disease, but in the absence of HTT, appears elevated when measured by HTRF^41^ (Fig 4e, Supplementary Fig. 5e, striatum, NTC vs. HTT, p< 0.0001; NTC vs. MSH3+HTT, p < 0.0001).

By 11 months, Q111 mice exhibited significantly increased aggregated HTT1a, with the highest accumulation in the striatum, followed by the cortex, and minimal aggregation in other brain regions (Supplementary Fig. 5f). This region-specific pattern corresponds with the areas where CAG expansion is most pronounced and where neuronal dysfunction occurs. Reducing Msh3 alone had no impact on HTT1a aggregation (Fig. 4f, Supplementary Fig. 5f striatum, NTC vs. MSH3, p= 0.4345), but co-reduction of Msh3 and HTT did significantly lower aggregated HTT1a (Fig. 4f, Supplementary Fig. 5f, striatum, NTC vs. MSH3+HTT, p < 0.01). Interestingly, reducing HTT alone showed an apparent increase in aggregated HTT1a (Fig. 4f, Supplementary Fig. 5f, striatum, NTC vs. HTT, p < 0.05). To confirm these results, we performed a western blot probing for HTT1a using antibody 1B12 on striatal lysates from a separate cohort of 12-month-old mice (same cohorts as mice in Fig. 1-3)^47^. Reducing Msh3 alone had no impact on aggregated HTT1a (Fig. 4g, Supplementary Fig. 6, NTC vs. MSH3, p = 0.2843), while co-reduction of Msh3 and Htt did reduce HTT1a aggregation (Fig. 4g, Supplementary Fig. 6, NTC vs. MSH3+HTT, p < 0.05). Importantly, reducing HTT alone did not increase 1B12 reactivity (Fig. 4g, Supplementary Fig. 6, NTC vs. HTT, p = 0.9997), suggesting that the increase observed in the HTRF assay (antibody 11G2) was relative rather than absolute.

In summary, HTT-lowering via siRNA reduced soluble full-length WT and full-length mutant HTT but not aggregated HTT1a. MSH3 knockdown prevented the loss of soluble full-length mHTT, suggesting expansion promotes aggregation. While MSH3 alone had no effect on HTT1a, combined MSH3 and HTT reduction decreased aggregated HTT1a levels across cohorts and assays. These results show that somatic expansion and HTT expression both drive HTT1a pathology, with MSH3 knockdown outperforming HTT-lowering alone and their combination offering enhanced therapeutic potential.

### Female Q111 mice display impaired nesting behavior compared to WT mice, a deficit that is rescued by MSH3 alone and or MSH3+HTT siRNA

We next sought to determine the effect of blocking somatic expansion of the CAG repeat in the *HTT* locus and lowering HTT expression on body weight and behavior in Q111 mice at 9, 11, and 12 months of age (Supplementary Fig. 7, 8a).

In general, male and female Q111 mice gained weight similar to male and female WT counterparts (Supplementary Fig. 7). Female Q111 mice showed no difference in weight regardless of their treatment group (Supplementary Fig. 7). Male Q111 mice treated with HTT and MSH3+HTT siRNA were slightly but statistically heavier than aCSF- or NTC-treated WT or Q111 mice (Supplementary Fig. 7).

At 9 and 11 months old, mice were assessed for overall motor gait, mobility, and function using motion sequencing (Supplementary Fig. 8b). There were no detectable differences between WT and Q111 mice observed at these ages, consistent with limited behavioral differences noted in Q111 in general and in particular at ages less than one year.

At 12 months of age to assess striatal motor and cognitive function, we performed a nesting assay that rates the quality of a nest build after 24 hours by a single mouse (1-5 scale, 1 is little nestlet manipulation and 5 is full nest observed). Female Q111 mice, but not male Q111 mice, were significantly worse at nesting than their WT counterparts (Supplementary Fig. 8c, Tukey’s multiple comparisons test, female WT aCSF vs female Q111 aCSF, p< 0.05, n=6,6). Nesting performance moderately rescued in Q111 mice treated with MSH3 siRNA, but not HTT siRNA (Supplementary Fig. 8c, Q111 NTC vs Q111 MSH3, p< 0.05). Nesting was markedly improved in Q111 cohorts that received MSH3+HTT siRNA (Supplementary Fig. 8c, Tukey’s multiple comparisons test, Q111 aCSF vs Q111 MSH3+HTT p < 0.01, Q111 NTC vs Q111 MSH3+HTT p < 0.001).

The remainder of the clasping and beam assays performed at 12 months old showed no measurable differences between WT and Q111 mice for male or female mice (Supplementary Fig. 8d, e, f).

### Blocking expansion combined with lowering HTT via siRNA is synergistic in rescuing neurodegenerative transcriptomic signatures in HD mice

The neurodegenerative transcriptomic signature is currently considered one of the gold standards for evaluating interventions in HD mouse models and characterizing disease severity in preclinical models^35^. To assess the neurodegenerative signature in mice from the experimental paradigm in Fig. 1, Supplementary Fig. 9a, (same cohorts as Fig. 1-3), we performed bulk mRNA-sequencing of the striatum. Principal component analysis of all reads did not find global batch effects or additional confounding factors in the dataset (Supplementary Fig. 9b,c). We then validated that the target genes (*Msh3* and/or *HTT*) were significantly lowered in the respective treatment groups (Supplementary Fig. 9d-o). RNA-sequencing analysis was performed using two sets of controls for completeness aCSF vehicle control and NTC siRNA (Fig. 5, Supplementary Fig. 9p,q). Our analysis confirmed that the dataset met quality control metrics, was technically valid for biological assessment, and achieved target knockdown in the expected groups. Importantly, the transcriptomic signatures of WT mice treated with divalent siRNA targeting *Msh3* or *HTT* confirm that siRNA-mediated Msh3 lowering and HTT lowering was safe and tolerated.

**Fig. 5.**
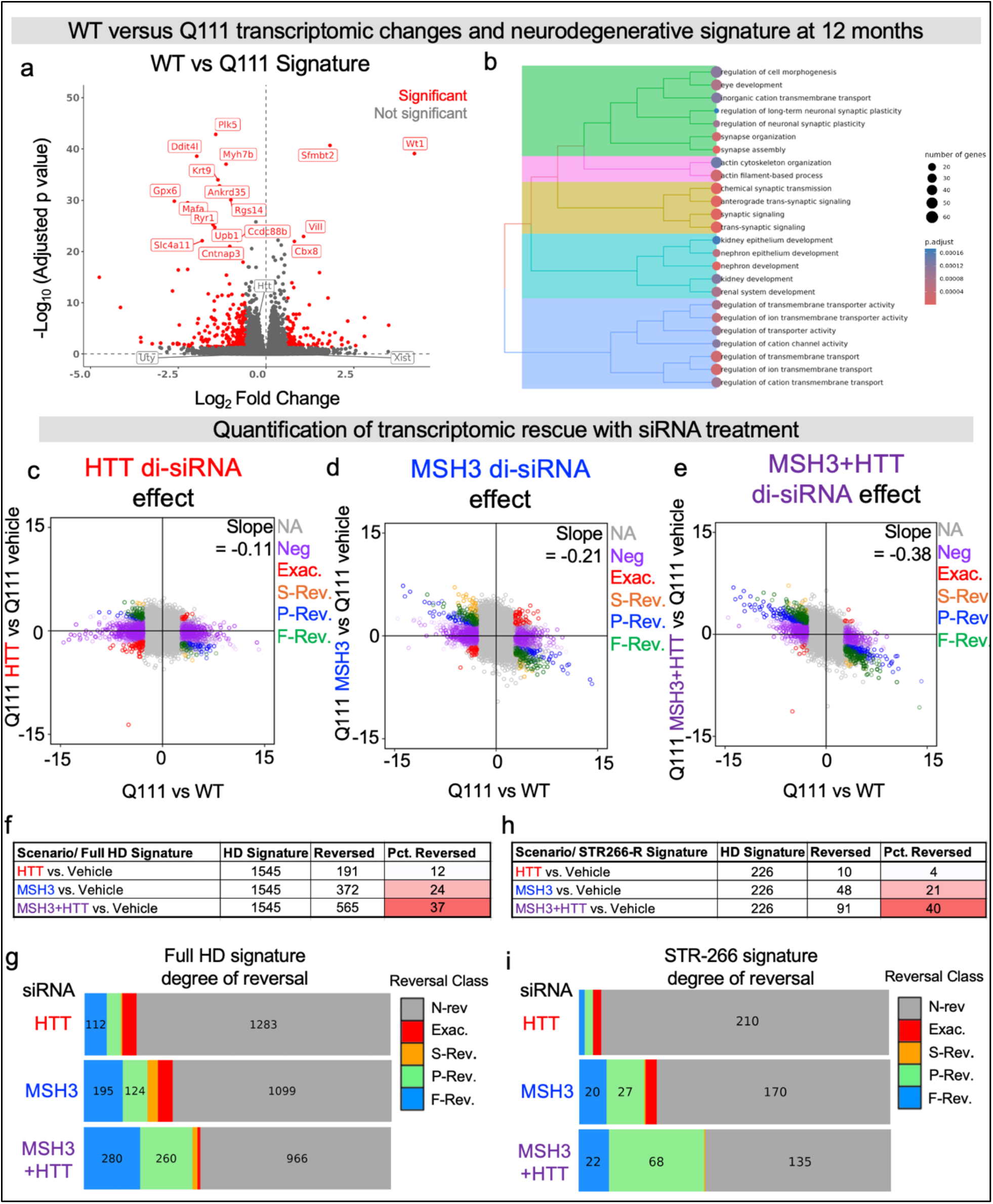
Bulk striatum transcriptomics demonstrates blocking CAG expansion with HTT co-silencing synergies to reverse HD neurodegenerative signature beyond either treatment alone. (**a**) Volcano plot of the differential gene expression analysis performed using DESeq2 between 12-month-old vehicle (aCSF-NTC) Q111 versus WT mice. Red indicates significant gene expression change. Significance determined by fold change > 1.5, FDR < 0.05. (**b**) Gene ontology biological processes (GOBP) pathway enrichment of the differentially expressed genes between vehicle Q111 and WT mice. (**c-e**) Overall gene reversal percentage of Q111 mice treated with (**c**) HTT siRNA (**d**) MSH3 siRNA or (**e**) MSH3+HTT siRNA combination compared to vehicle Q111 mice. Color indicates the degree of reversal classified as full, partial, super, exacerbation, or negligible. (**f-i**) Prevention, reversal, recuse (PuRR) classification of the degree of gene expression reversal. Number of genes differentially expressed or reversed by siRNA treatment in the (**f**) whole HD signature or (**h**) select STR266 HD signature genes across treatment cohorts as defined by Marchionini et al. 2022 ^48^. (**g-i**) classification of the degree of reversal for the (**g**) whole HD signature or (**i**) STR266 top gene list. Degree of gene expression reversal color coded as: Blue-full reversal (F-Rev); green-partial reversal (P-Rev); orange-super reversal (S-Rev); red-exacerbation (Exac); gray-negligible reversal (N-Rev). Transcriptomics performed on the same mouse cohorts presente**d in Figures 1-3**. N=9-12 mice per treatment group.

To understand which pathways were represented, and to what extent, in the disease signature in Q111 mice, we examined differentially expressed genes (DEGs) in control WT versus control Q111 mice. (Fig. 5a). The disease signature was defined as genes differentially expressed in both the aCSF and NTC Q111 cohorts (vehicle signature), ensuring robustness given that two controls were used throughout the experiment. The Q111 signature showed profound gene expression changes compared to WT mice (1545 DEGs) (Fig. 5a,b). These DEGs align with known pathways dysregulated in HD, primarily neuronal dysfunction, synaptic dysfunction, and proteomic stress (Fig. 5b).

We next analyzed the HD reversal score^48^, a metric for assessing disease reversal in HD models. The reversal fraction quantifies gene changes between WT and Q111 mice and estimates the extent of return to WT levels with treatment^48^. The degree of reversal is classified as Expression_treatment_= α*Expression_disease_. There are five categories of gene reversal: super reversal (α < –1.3); full reversal (–1.3 < α < –0.7); partial reversal (–0.7 < α < –0.3); negligible reversal (– 0.3 < α < 0.3); and exacerbation (α > 0.3)^48^.

Overall, lowering HTT alone had little impact on gene rescue (Fig. 5c), but lowering Msh3 alone or co-lowering both significantly reversed the disease signature (Fig. 5d, e). To better quantify disease reversal, we measured the number and types of genes reversed by divalent siRNA treatment, first by looking at the treatment impact on all DEGs (HD Signature) (Fig. 5f,g, Supplementary Fig. 10a). Compared to vehicle treatment, HTT lowering led to a reversal of 191 out of 1545 DEGs (12% reversed) (Fig. 5f,g). MSH3 lowering reversed 372 out of 1545 DEGs (24% reversed) and reducing both MSH3 and HTT reversed 565 out of 1545 DEGs (37% reversed) (Fig. 5f,g).

We next analyzed the STR266 signature, a set of 266 genes consistently dysregulated across HD mouse models in the striatum, representing a specific and widely applicable HD signature^49^. The vehicle Q111 signature in this study had 226 out of 266 STR266 genes dysregulated. Compared to the vehicle Q111 signature, HTT lowering led to a reversal of 10out of 226 DEGs (4% reversed) (Fig. 5h,i). MSH3 lowering reversed 48 out of 226 DEGs (21% reversed), and lowering both targets reversed 91 out of 226 DEGs (40% reversed) (Fig. 5h,i, Supplementary Fig. 9b).

In summary, co-reduction of MSH3 and HTT showed substantial transcriptional rescue, suggesting an additive or even synergistic effect on disease-associated pathways. These transcriptomic findings align with our results at the protein level, which showed a less progressed mHTT phenotype and a significant reduction in HTT1a aggregation following the co-reduction of MSH3 and HTT. MSH3 lowering alone is also sufficient to reduce mHTT aggregation and drive transcriptional rescue. A time-resolved HD model for each treatment condition is presented in Fig. 6. Collectively, this study supports a disease-modifying treatment for HD.

**Fig. 6:**
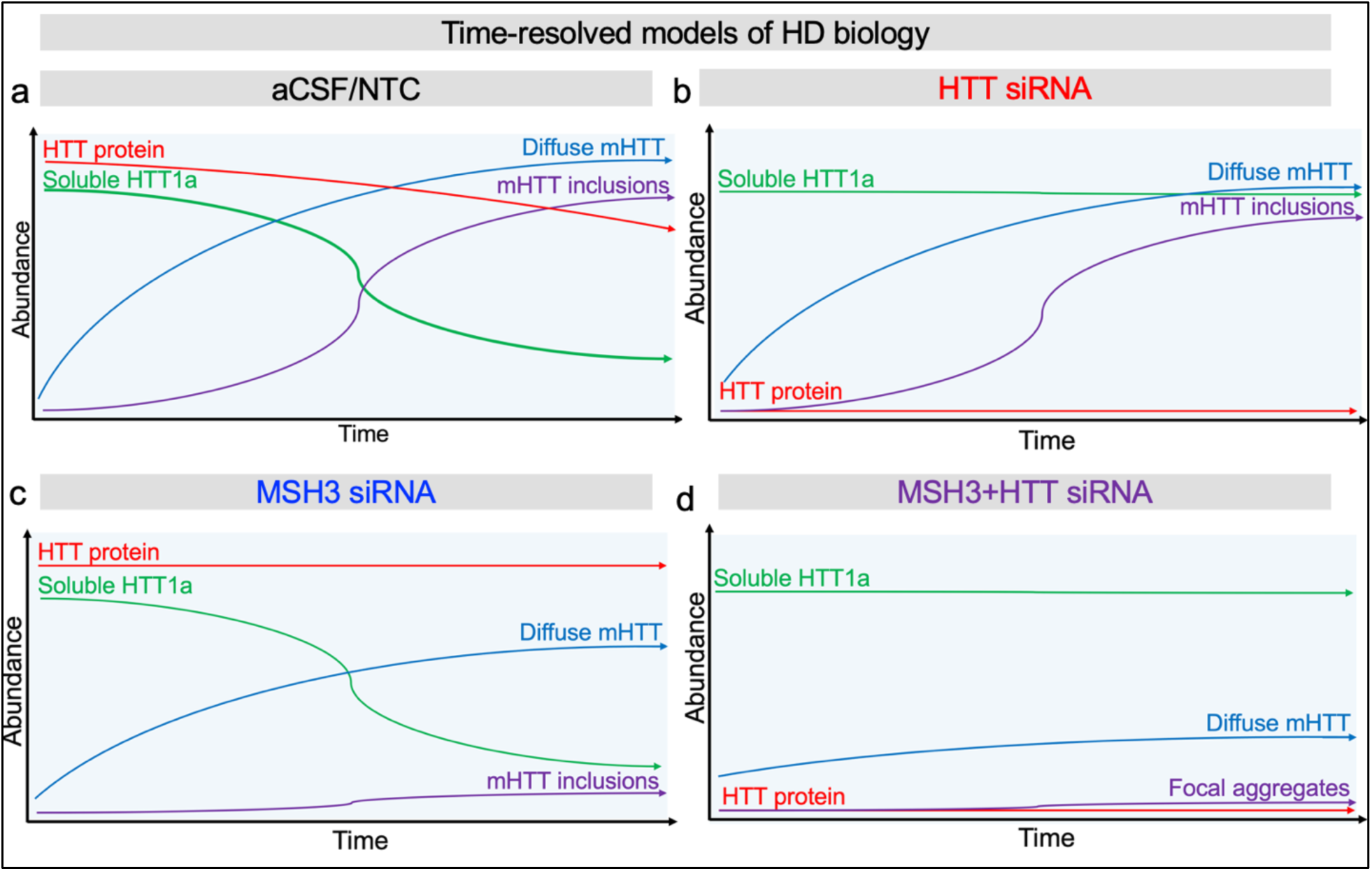
Time-resolved HTT, HTT1a, CAG expansion model of HD biology. Model of HTT species across time with (**a**) artificial CSF or non-targeting control siRNA, (**b**) HTT siRNA, (**c**) MSH3 siRNA, (**d**) MSH3+HTT siRNA combination. Total full-length mutant HTT (red), soluble HTT1a (green), diffuse mHTT aggregates (blue), or mHTT inclusions (purple).

## DISCUSSION

We report that blocking somatic expansion with therapeutic-grade siRNA can modify molecular and behavioral phenotypes in a mouse model of HD, by reducing focal mHTT aggregation, shifting mHTT protein status to an earlier disease state, and reversing transcriptional signatures of neurodegeneration in HD. Blocking somatic CAG expansion in combination with HTT lowering further slows mHTT aggregation, rescues disease signatures, and reduces HTT1a aggregate levels. These effects were observed in the context of at least 110-120 CAG repeats. Lowering HTT alone had little impact on mHTT aggregation or transcriptional rescue. Overall, this work identifies a disease-modifying siRNA combination therapy for testing in testing in people with HD.

Consistent with our findings in Q111 mice, blocking CAG expansion via genetic knockout of *Msh3* prevented mHTT aggregation and strongly reversed HD signatures in the Q140 (140-150 CAGs) mouse model^22^. However, *Msh3* knockout had little to no effect on these outcomes in the Q175 (185-190 CAGs) mouse model. This suggests that blocking expansion is mainly disease-modifying when performed at CAG lengths under 140 repeats^15, 22, 25^, supporting the threshold model of HD^8, 15, 22, 25, 26^ and the associated “therapeutic” range of 40–150 CAG repeats in the CNS. How this translates to human patient age is unclear, but pre-symptomatic HD progression biomarkers including blood CAG lengths, neuroimaging, and neurofilament light levels can be used to guide the design of clinical trials with this question in mind, of which extensive work is already underway^14, 50–52^.

Multiple MSH3-lowering approaches are in preclinical development, including antisense oligonucleotides (ASOs)^53^, small molecules, and viral vectors. Somatic expansion can also be targeted by modulating other MMR pathway proteins or directly binding the HTT locus^54, 55^. Collectively, these approaches represent a diverse and growing toolkit to slow or prevent disease progression in HD by addressing somatic expansion. Notably, the divalent siRNA used in this study achieves potent, CNS-specific knockdown with a single injection, sustaining its effect for 6–9 months in large animals. Among potential targets, MSH3 stands out for its profound impact on CAG expansion, its exceptional safety profile, and minimal transcriptomic impact in wild-type mice, supporting its role as a precise, durable, and disease-specific therapeutic strategy that preserves broader MMR and cellular functions.

In the present study, we found that lowering HTT had a minimal impact on mHTT aggregate counts and transcriptomic rescue when somatic expansion is ongoing. Yet, combined HTT lowering and blocked somatic expansion had a synergistic impact on reducing aggregates and transcriptomic rescue, emphasizing that HTT-lowering therapies are most effective when CAG repeat lengths are concurrently stabilized. The chemically stabilized divalent siRNA targeting *HTT* or *MSH3* in the present study are both clinical trial-ready. Additional clinically advanced modalities including ASOs^53^, lipophilic conjugated siRNA (C16),^56^ and transferrin receptor-coupled oligonucleotides that can cross the blood-brain-barrier^57^ can also be programmed to silence *MSH3* alone or in combination with *HTT*.

In advanced HD (i.e., cases of extremely long CAG repeat tracts), siRNA-mediated lowering of HTT is less effective, likely because siRNA cannot efficiently target aggregated mRNA^58^. Nuclear aggregates of *HTT* mRNA are more likely to have ultralong CAG lengths^15^. Beyond controlling *HTT* CAG lengths, therapeutic approaches (e.g., siRNA, ASOs, AAVs) that silence HTT1a to break down aggregates are ongoing^59, 60^. Early evidence from these studies suggests that targeting HTT1a is more effective at preventing mHTT aggregation than targeting full-length HTT^60^. Future work characterizing the interplay between somatic expansion and HTT1a is essential to map the potential treatment landscape for HD.

Aggregation is also influenced by HTT concentration—only when both repeat length and HTT levels are low does aggregation decrease significantly. Blocking expansion alone did not prevent microaggregates in older mice, despite similar CAG lengths to younger mice without aggregates. This suggests that mHTT aggregation is time-dependent and driven by ongoing HTT transcription, highlighting a temporal component in HD progression beyond somatic expansion.

Therapeutic modulation of MSH3 and HTT appears safe for at least one year in WT and Q111 mice, with minimal transcriptional changes. Divalent siRNAs would likely be administered intracerebroventricularly, intraparenchymally, or intrathecally. Intrathecal injection is used for the FDA-approved ASO drug, Nusinersen^29^. The use of intraventricular and intraparenchymal administrations is still emerging, but one might expect treatment intervals of 6–9 months to maintain efficacy. Any nucleic acid material injected into the cerebrospinal fluid (CSF) or parenchyma ultimately clears through the liver, raising the possibility of residual gene silencing in this tissue. Aptamers that bind systemically available siRNA to block their activity have been developed, ensuring CNS-specific gene modulation^61^.

MSH3 remains an attractive target due to its specific role in DNA repair. Knockout studies in mice indicate that MSH3 deficiency alone does not lead to systemic cancer development over one year^62^. In humans, certain MSH3 polymorphisms are associated with an increased risk of proximal colon cancer, though this risk is less pronounced than that associated with MSH2 mutations^63^. Consequently, any future MMR-targeted therapeutic may require routine colon cancer screening throughout the treatment course to manage any new cancer risk imposed. Prior work suggests divalent siRNA therapeutics are primarily cleared by the liver further mitigating an effect of MSH3 lowering in the colon^28, 61^. Nevertheless, long-term safety studies in large animals are required.

Identifying disease-modifying treatments for HD is challenging due to limitations in mouse models, including a lag between biochemical changes and behavioral phenotypes, which may take 15–18 months to appear^32, 33^. To assess early disease drivers, we evaluated mice at 12 months, identifying a nesting deficit in female Q111 mice as the primary behavioral phenotype—fully rescued by MSH3 or MSH3+HTT siRNA. As behavioral readouts are limited, transcriptomic rescue is emerging as a key surrogate for neurodegeneration reversal in HD models^22, 25, 35^.

A challenge in this study was choosing the most accurate control (aCSF vs. NTC siRNA). We used a shared transcriptomic signature from both to define HD-associated changes, aligning with known dysregulated pathways, though defining a definitive HD signature in mice remains difficult^22, 25, 35, 48^. In clinical trials, natural history data replace oligonucleotide-based controls, highlighting a limitation of preclinical models.

Unlike humans with somatic mosaicism, all cells in Q111 mice carry expanded CAG repeats, accelerating disease and complicating assessments of cell-specific vulnerability. However, this may mean the benefits of blocking somatic expansion are even greater in humans. Our findings support that eliminating mHTT with long CAG tracts is crucial for slowing HD, and while we used pan-allele HTT silencing, other more selective approaches may offer additional therapeutic promise.

This work demonstrates that blocking somatic expansion combined with HTT-lowering is a clinically promising therapeutic strategy for HD. While the biological rationale is straightforward, HD clinical trials present unique challenges. A key consideration is patient identification, as much of the disease burden may still be modifiable through interventions that block disease progression, particularly when applied early, before symptom onset, rather than later in symptomatic stages. Clinical trials for HD must accommodate the slow progression and heterogeneity of the disease, and identifying registerable outcomes measurable over 2–5 years is essential to measure treatment efficacy^64, 65^. Additionally, assessing the on-target efficacy of approaches that block somatic expansion remains an unresolved challenge. Efforts to validate biomarkers for CAG repeat stability (across CAG expansion disorders) and mHTT reduction are ongoing^66, 67^, although the utility of blood-based biomarkers may be limited due to the CNS-specific activity of divalent siRNA. Despite this, significant efforts are underway in the HD biomarker field. Neurofilament light chain (NfL) remains a critical biomarker of neuronal damage, offering insights into disease onset, progression, and treatment toxicity^52, 68–70^. Overall, clinical biomarker validation is essential for translating these preclinical findings into effective treatments for humans. In addition, progress is being made to develop an mHTT positron emission tomography (PET) ligand that detects aggregated mHTT in the brain^71, 72^. PET ligands may become an important tool clinically to monitor the effects of therapeutics including MSH3 and HTT agents predicted to affect mHTT levels and aggregation.

In conclusion, blocking somatic expansion, achievable with interventional divalent siRNA, is key to slowing HD progression. Combined with HTT-lowering, it synergistically eliminates mHTT aggregates, reduces HTT1a, improves nesting behavior, and reverses neurodegenerative gene signatures in mice, guiding next-generation HD therapies.

## MATERIALS AND METHODS

### Oligonucleotide synthesis

Solid-phase synthesis with phosphoramidite chemistry was performed using a MerMade12 automated synthesizer (Biosearch Technologies, Novato, CA). Phosphoramidites used were 2’OMe and 2’F with standard protecting groups. 5’-(E)-Vinyl tetraphosphonate (pivaloyloxymethyl) 2’-O-methyl-uridine 3’-CE was used for the addition of the 5’-(E)-Vinyl phosphonate. Phosphoramidites were brought into solution at 0.10M using anhydrous acetonitrile (ACN). 2’-OMe-uridine was an exception and was brought into solution at 0.10M with anhydrous ACN and 15% dimethylformamide. On-column detritylation was done using 3% trichloroacetic acid in dichloromethane. The activator was 0.25M 5-(Benzylthio)-1H-tetrazole (BTT) with 4-minute coupling times. CAP A (20% N-methylimidazole in ACN) and CAP B (20% acetic anhydride and 30% 2,6-lutidine in ACN) were the capping reagents used. Oxidation or sulfurization was performed using 0.05M iodine in pyridine and water (9:1, v/v) or 0.10M 3-((dimethylaminomethylene)amino)-3H-1,2,4-dithiazole-5-thione in pyridine (DDTT) for 3 minutes. All phosphoramidites were purchased from Hongene Biotech (Union City, CA) and Chemgenes (Wilmington, MA), and all reagents were purchased from Chemgenes. The solid-support controlled pore glass used to grow the oligonucleotides included a long-chain alkyl amine (LCAA) and was functionalized with a succinyl linker with 500Ä Unylinker terminus (Chemgenes) or a custom-made 500Ä di-trityl protected support connected by tetraethylene glycol linker (Hongene).

### Oligonucleotide cleavage, deprotection and purification

All oligonucleotides underwent cleavage from the solid support and removal of protecting groups. Oligonucleotides containing 5’-(E)-Vinyl phosphonate were cleaved and deprotected with 3% diethylamine in ammonium hydroxide for 20h, 35°C with slight agitation. The divalent oligonucleotides were cleaved and deprotected using 1:1 40% aqueous monomethylamine with ammonium hydroxide for 2h, 25°C with slight agitation. The oligonucleotides were then filtered and rinsed with 5% ACN in water and dried overnight by centrifugal vacuum concentration. Purification was performed with Source 15Q anion exchange resin on an Agilent 1290 Infinity II HPLC system. A linear gradient of 30 – 70% in 30 minutes at 50°C was used, and peaks were monitored at 260nm for all oligonucleotides. Select fractions were combined and desalted with size exclusion chromatography with Sephadex G-25 on an Akta FPLC. Oligonucleotide purity and identity were confirmed using IP-RP on an Agilent 6530 Accurate-mass Q-TOF.

### Divalent siRNA duplexing and dosing

In vivo grade siRNA was duplexed to 10 nmol divalent scaffold/10 μL (2000 nM). To eliminate acute neurotoxicity, divalent siRNA was prepared in a calcium- and magnesium-enriched artificial CSF (aCSF) buffer ^36^. Vehicle control was the same calcium- and magnesium-enriched aCSF. The combination siRNA mixture cohort received 10 nmol Msh3-targeting sequence and 10 nmol HTT-targeting sequence each, in 10 μL total.

### Mouse work

All mouse experiments were performed in alignment with the Institutional Animal Care and Use Committee at the University of Massachusetts Chan Medical School protocol 202000010. All mice were bred and born at Jackson Labs in Farmington, CT. Beginning at 6 weeks old, mice were transferred to the pathogen-free UMass Chan animal facilities where they were housed and monitored with a 12h light:12 hr dark cycle at 23 ± 1°C and 50 ± 20% humidity. Mice were allowed free access to food and water. Mice used were either wild-type C57/B6 Wild-type or Hdh^Q111^ Het mice, JAX Strain ID 370624, Mc_Q111 KI; C57BL/6J background, heterozygous.

### Stereotaxic intracerebroventricular injections

Mice were anesthetized using isoflurane. Once fully anesthetized, the head was shaved, and a small skin incision was made to reveal the scalp. A burr hole was created using coordinates from bregma, mediolateral +- 1mm, posterior -0.2mm, and ventral -2.5mm. Five μL per ventricle was injected at a rate of 750 nL/min. This process was repeated bilaterally. Post-procedure, mice were given Meloxicam ER and saline. Mice were monitored until sternal, daily for 72 hours post produce, and weekly for the duration of the experiment.

### Behavior: nesting, clasping, beam walking

Motion sequencing was performed according to parameters validated by the assay developer^73^. Motion sequence data was analyzed as a correction across genotype vector with 9-month-old aCSF-treated WT and Q111 cohorts used to form a “behavior progression” vector. Treatment cohorts are assessed by where they fall on the defined vector. Nesting assay: mice were singly housed, and enrichment was removed and replaced with one nestlet overnight. The next morning, nest building was assessed based on the nesting protocol described previously ^74^: 5 is full nest, 1 untouched nestlet. Hindlimb clasping: mice were held midway along tail for 30 seconds. Mice received a score of 0= no clasping events, 1= one limb retracted briefly, 2 = one limb retracted fully 3= bilateral clasping observed^75^. Beam Walking: mice were trained on 4-foot long, a 1-inch diameter beam for 3 sessions over 1 week. To assess beam walking performance, mice were placed on beam. Each mouse performed 3 consecutive walks end-to-end. Videos were recorded from two perspectives and analyzed for foot slips, time, and prods by a third party without information about the genotype or treatment group.

### mRNA quantification

At the time of sacrifice, 1.5 mm x 1 mm round punches across brain regions were collected and stored in RNAlater overnight. One punch was lysed into 600 μL homogenizing mix (Invitrogen ThermoFisher QG0517). mRNA was measured using the branched DNA Quantigene Singleplex assay (Invitrogen ThermoFisher QS0016) probing for *Msh3* (Mouse, SA-3030208), *Htt* (Mouse, SB-14150), or *Hprt* (Mouse, SB-15463) housekeeping levels. mRNA levels were first normalized to housekeeping controls, and treatment efficacy was quantified relative to non-targeting control (NTC) levels.

### Protein preparation, western blot analysis, and quantification

Crude homogenates were prepared by homogenizing frozen tissue punches on ice in 10mM HEPES pH7.2, 250mM sucrose, 1mM EDTA, + protease inhibitor tablet (Roche) + 1mM NaF + 1mM Na3VO4 then sonicating for 10 seconds. Protein concentration was determined using the Bradford method (BioRad), and 10-20 µg of protein were separated by SDS-PAGE on 3-8% Tris-acetate Criterion gels (BioRad). Protein was transferred to nitrocellulose using a TransBlot Turbo apparatus (BioRad). Western blots were performed as previously described (Sapp et al., 2020) using antibodies to HTT (aa1-17, 1:2000, DiFiglia et al., 1995), Msh3 (1:500, Santa Cruz sc-271079), HTT1a (5ug/ml, P90 1B12, Coriell Institute), Gapdh (1:10000, MilliporeSigma), and B-actin (1:5000, MilliporeSigma). Images were acquired using ChemiDoc XRS+, and pixel intensity quantification was performed using ImageJ software. The bands for HTT, Msh3, Gapdh, and B-actin were manually circled, and total signal intensity was determined by multiplying the area by the average signal intensity. For HTT1a, (Antibody 1B12) the average signal intensity in equal areas from the top of the smear to the band at ∼75kD was determined for each sample. The total or average signal intensities were normalized to Gapdh or B-actin loading controls. Statistical analyses were One-way ANOVA with Tukey’s multiple comparison tests.

### Fragment analysis PCR

A 1.5-mm x 1-mm round striatum or medial cortex punch was flash-frozen at the time of takedown. DNA extraction was performed using Qiagen DNeasy Blood and Tissue Kit (Qiagen Cat. 69506). Fragment analysis PCR was performed using the AmpliTaq 360 DNA Polymerase kit (ThermoFisher Cat. 4398818). 150 ng of DNA per sample was loaded into the PCR reaction. Forward primer CAG1:6FAM-ATG AAG GCC TTC GAG TCC CTC AAG TCC TTC. Reverse primer HU3: GGC GGC TGA GGA AGC TGA GGA. Thermocycling performed at: 94C, 90s; [94C, 30s; 63.4C, 30s; 72C, 90s]x 35, 72C, 10m. Fragment analysis was performed at the University of Arizona Genetics Core. Somatic instability was visualized with the ThermoFischer PeakScanner software and quantified using the index developed by Lee et al. ^37^.

### Immunohistochemistry/Immunofluorescence

Half mouse brains were drop-fixed in 10% neutralized buffered formalin (X). 48 hours later, fixed brains were moved to phosphate-buffered saline. Mouse hemi sections were sliced using a vibratome into 40-μM thick slices. For long-term storage, brains were stored in PBS with sodium azide. For immunohistochemistry, tissue was permeabilized with 3% H_2_O_2_ for 3 minutes, followed by 0.1% Triton-X-100 at RT for 20 minutes. Tissue was blocked with the M.O.M. Mouse Ig Blocking Reagent using Vector M.O.M. kit with ImmPRESS Peroxidase Polymer (cat. # MP-2400) for 1 hour and 2.5% normal house serum for 5 minutes. Primary antibody EM48 Antibody (MAB5374, 1:100 in 2.5% normal horse serum) was applied overnight at 4°C. Secondary antibody Vector M.O.M. ImmPRESS Reagent (anti-mouse IgG) was applied at 10 minutes RT. Sections were stained with 1X Metal Enhanced DAB Substrate Kit (ThermoFisher, cat. #34065) and mounted onto coverslips.

### Image acquisition and quantification

Images were collected with an Andor Dragonfly 505 on a Leica inverted DMi8 using confocal mode equipped with a Leica HC PL APO 40x/1.40 OIL CS2 objective using transmitted light. Images were acquired with an Andor Zyla sCMOS 4.2 Plus camera controlled with Fusion (2.4.0.14) software. Brightness and contrast were adjusted using Fiji3 (2.0.0-rc-69/1.52p). Three consecutive striatal slices (40 µM thick) were imaged for each mouse. Within each striation slice, the center of each stratum was imaged followed but the field of view immediately above and below at 40x. This was repeated for 2 more striatal slices for a total of nine images per mouse acquired. Four mice per group were imaged. Quantification was performed without information on the genotype or treatment group. EM48 staining was characterized as focal or diffuse during quantification.

### Homogenous time resolved fluorescence (HTRF) Protein lysate preparation

For individual Q111 mouse brain tissue lysate preparation, a 10% (w/v) total protein homogenate was prepared in ice-cold bioassay buffer (phosphate-buffered saline (PBS), 1% Triton-X-100) with complete protease inhibitor cocktail tablets (Roche), by homogenizing three times for 30 s in Lysing matrix D tubes at 6.5 m/s (MP Biomedicals) in a Fast-Prep-24^TM^ instrument (MP Biomedicals), and up to 10 μl lysate was aliquoted per well in triplicate for bioassay analysis. For aggregation assays, 10 μl crude lysate was used, whereas, for soluble assays, 10 μl of supernatant was used after brief centrifugation at 3500xg for 10 min.

### HTRF assay

Q111 mouse brain homogenates to a final volume of 10 μl were pipetted in triplicate into a 384-well (pure white, low volume, conical) proxiplate (Greiner Bio-One). Lysate dilutions and antibody concentrations are summarized in Supplementary Table 2. For HTRF assays, terbium cryptate donor and either d2 or alexa-488 acceptor antibodies were added per well in 5 μL HTRF detection buffer (50 mM NaH_2_PO_4_, 0.2 M KF, 0.1% bovine serum albumin (BSA), 0.05% Tween-20) with cOmplete protease inhibitor cocktail tablets (Roche). Plates were incubated for 3 h on an orbital shaker (250 rpm) at room temperature before the 15 μL reaction was read using an EnVision (Revvity) plate reader. For d2 acceptor detection, HTRF parameters were mirror: LANCE/DELFIA 412, excitation filter: UV2 (TRF) 320 (111), emission filter: APC 665 (205), 2nd emission filter: Europium 615 (203), excitation: 100%, delay: 60μs, window time: 150μs, number of flashes: 100, 2nd detector flashes: 100, time between flashes: 2000μs. For Alexa-488 acceptor detection, HTRF parameters were mirror: LANCE/DELFIA 412, excitation filter: UV2 (TRF) 320 (111), emission filter: TRF 520 (275), 2nd emission filter: TRF 495 (276), excitation: 100%, delay: 150μs, window time: 800μs, number of flashes: 100, 2nd detector flashes: 100, time between flashes: 2000μs.

### RNA-sequencing and Transcriptomic Analysis

1.5-mm x 1-mm striatum punches were flash-frozen at the time of sacrifice. RNA from one punch per mouse was extracted and purified using the Monarch Total RNA Miniprep (T2010S) kit. RNA QC was verified using the RNA ScreenTape (Agilent #5067). Purified RNA was sent to Genewiz for bulk RNA sequencing. Reads were aligned with Kallisto^76^ against the mouse reference genome, mm10 or the Qiagen Omicsoft alignment workflow against the mouse reference genome GRCm38. Quality control and analysis were performed at Rancho Biosciences. The differential gene expression analysis and transcriptomic rescue analysis used the posterior probability-based (PP) method developed Marchionini et al 2022^48^. This approach gives a reversal probability (RP) to each gene. A gene list of differentially expressed genes is created between WT and Q111 mice based on genes that show a fold change of at least 15% and adjusted *P* < 0.05 after multiple-test correction. The impact of treatment is classified as Expression_treatment_= α*Expression_disease._ The degree of reversal for each gene is classified as classified by its α. There are 5 categories of gene reversal: α < –1.3 (super-reversal); –1.3 < α < –0.7 (full reversal): –0.7 < α < –0.3 (partial reversal): –0.3 < α < 0.3 (negligible reversal) or α > 0.3 (exacerbation). To generate the HD signature for this analysis, differential expression analysis was performed comparing Q111 aCSF and NTC samples to WT aCSF and NTC samples using the treatment as a covariate to provide a broad HD signature representing both controls. Differential expression tests were performed in R using the DESeq2^77^ package with independent filtering disabled, and a significance threshold of an adjusted P < 0.05 after multiple-test correction. The ‘HD signature’ dysregulated gene list was tested for gene set over-representation against the GOBP (gene ontology biological process)^78^ version 18.9.29 using the enricher function within the R clusterProfiler^79^ package. A clusterProfiler q < 0.05 was deemed significant.

### Statistics

Statistics were performed using GraphPad Prism Version 10.2.3. When comparing two groups, a student’s t-test was used. When comparing more than two groups with one variable, a one-way ANOVA with multiple comparisons was used. When comparing two variables, a two-way ANOVA with multiple comparisons was used. Within two-way ANOVAs, Dunnett’s multiple comparisons test was performed when comparing the treatment effect the brain (regardless of brain region) or direct comparisons against NTC. Tukey’s multiple comparisons test was used when comparing treatment effect within specific brain regions. When analyzing aggregate abundance, one-way ANOVA with Holm-Sidak correction test was used. On behavioral score-based assays, one-way ANOVA with Dunn’s multiple comparisons test was performed. In vivo experiments were performed with N=6-12 mice per condition, and immunohistochemistry or immunofluorescence analyzed N=4 mice per treatment group. Immunohistochemistry analysis and behavior analysis were performed blinded to mouse genotype and treatment group. Where graphically reported, statistics are denoted as, * p < 0.05, ** p < 0.01, *** p < 0.001, **** p < 0.0001.

## Acknowledgments

The authors thank Emily Hablerlin for reviewing the manuscript and providing feedback. The authors thank the CHDI Foundation (A-5038 to AK, NA, MD; A-19405 to AK; A-19252 to GB). AK was supported by NIH R01 NS104022, S10 OD020012, S10 OD036329. JNB was supported by NINDS F31 NS132424. PLG was supported by Smith Family Foundation Odyssey Award and NIH NIA R56AG089801.

