## Supplemental Data for "Blocking somatic repeat expansion and lowering huntingtin via RNA interference synergize to prevent Huntington’s disease pathogenesis in mice"

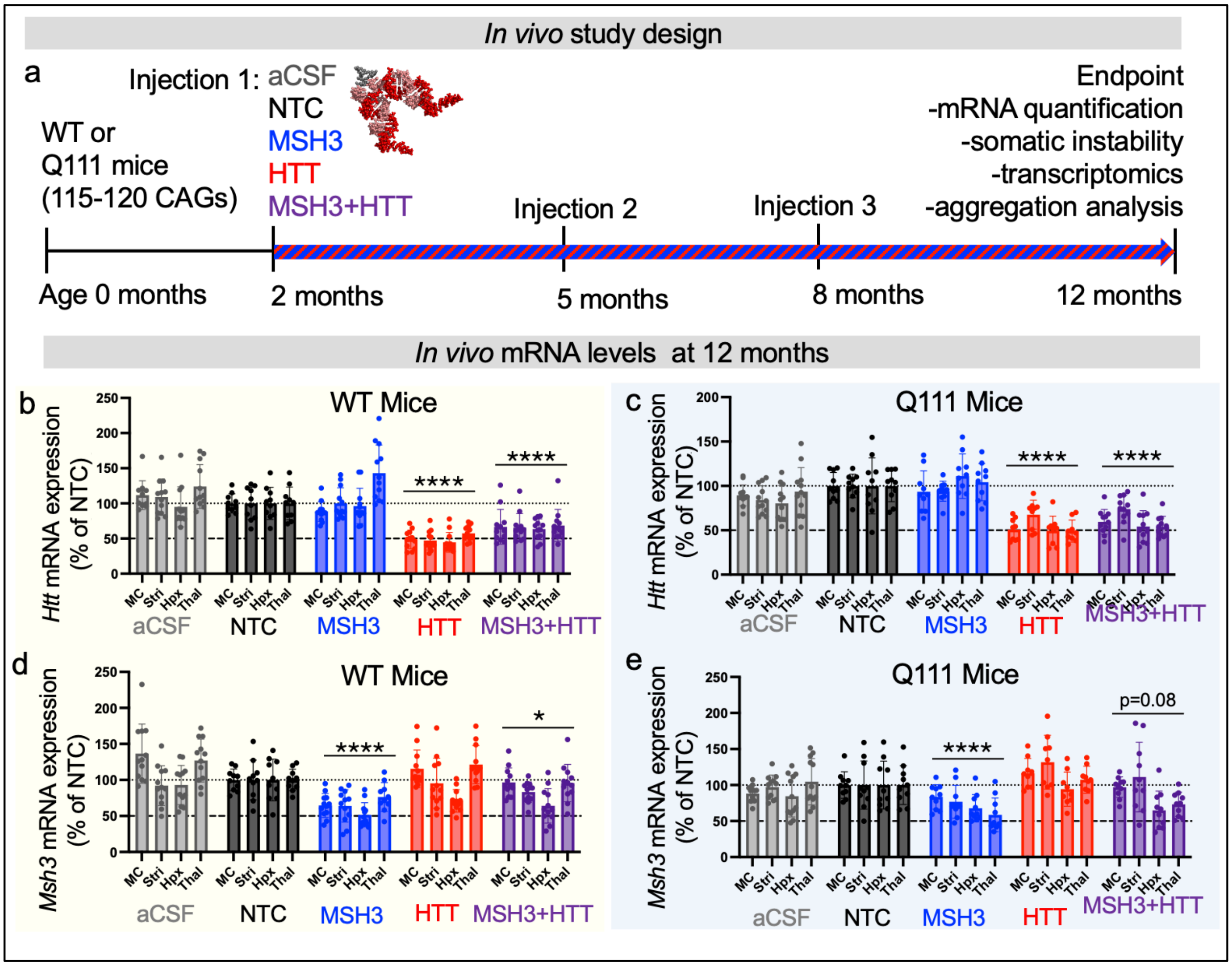


**Supplementary Fig. 1. mRNA quantified in 12-month-old WT or Q111 cohorts at experimental endpoint.** (**a**) Experimental design (**b**) *Htt* mRNA in WT mice, (**c**) *Htt* mRNA in Q111 mice, (**d**) *Msh3* mRNA in WT mice, (**e**) *Msh3* in Q111 mice. MC, medial cortex; Stri, striatum; Hpx, hippocampus; Thal, thalamus. mRNA quantified using Quantigene branched DNA assay.

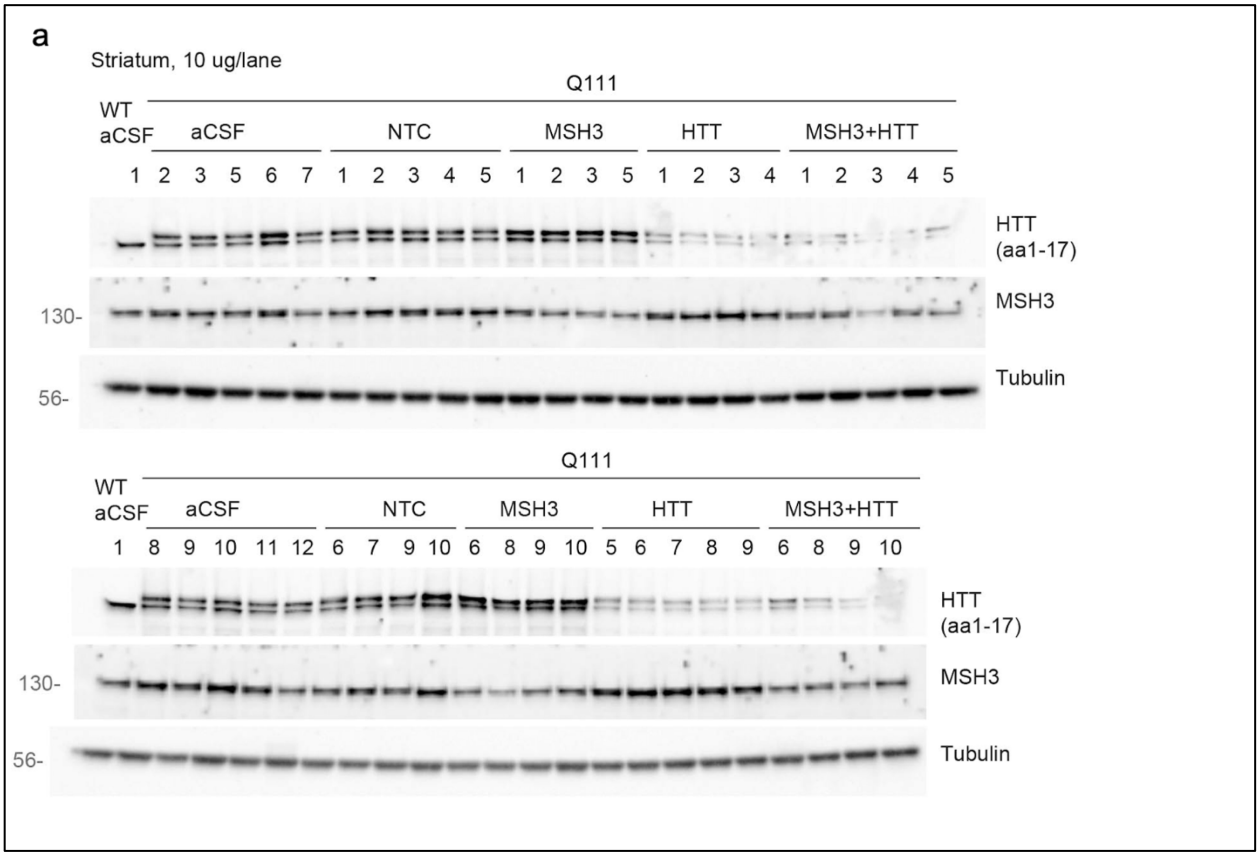


**Supplementary Fig. 2. Western blots used to quantify MSH3 and HTT protein levels.** Striatum tissue, Q111 mice aged 12 months. Each lane is lysate from a single WT mouse treated with aCSF or Q111 mouse treated with aCSF or divalent siRNA programmed with sequences targeting non-targeting control (NTC), MSH3, HTT, or both MSH3 and HTT. HTT detected using anti-HTT antibody Ab1 (aa1–17)[80].

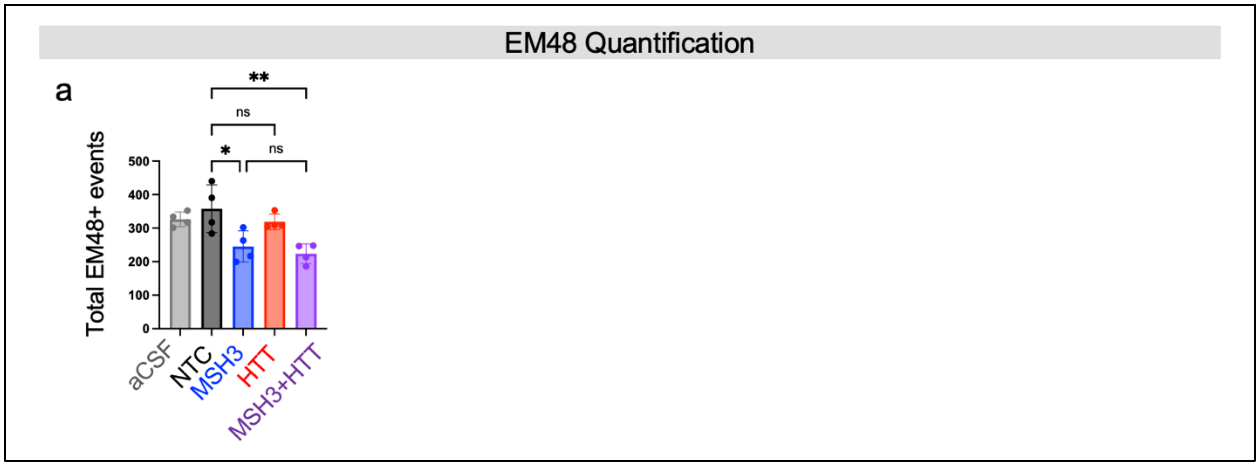


**Supplementary Fig. 3. Total EM48+ mHTT event across treatment cohorts.** Total EM48+ event is defined as the sum of any EM48+ diffuse or puncta event per region of interest.

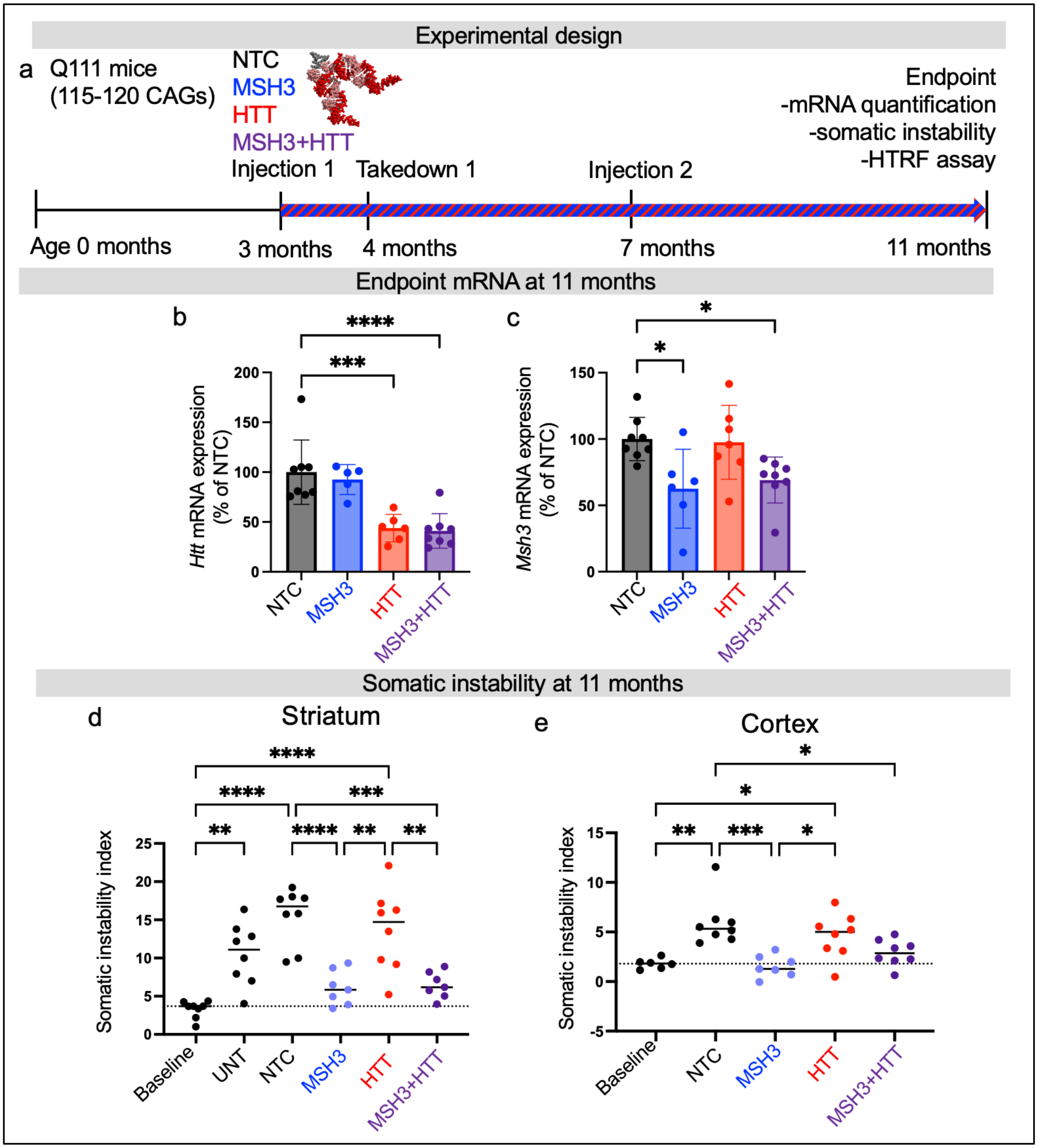


**Supplementary Fig. 4. MSH3 and HTT knockdown achieved; expansion blocked in expected siRNA cohorts used to assess HTT species abundance.** (**a**) Experimental paradigm of WT and Q111 cohorts used to assess HTT species using with homogenous time-resolved fluorescence. (**b,c**) mRNA levels at experimental endpoints (11 months) probing (**b**) *Htt* or (**d**) *Msh3*. mRNA quantified using Quantigene branched DNA assay. (d,e) somatic instability index in (**d**) striatum and (**e**) medial cortex at endpoint (11 months). Baseline is 3-month-old untreated Q111 striatum. UNT, untreated, non-targeting control (NTC), siRNA against MSH3, HTT, or MSH3+HTT combination at 11 months.

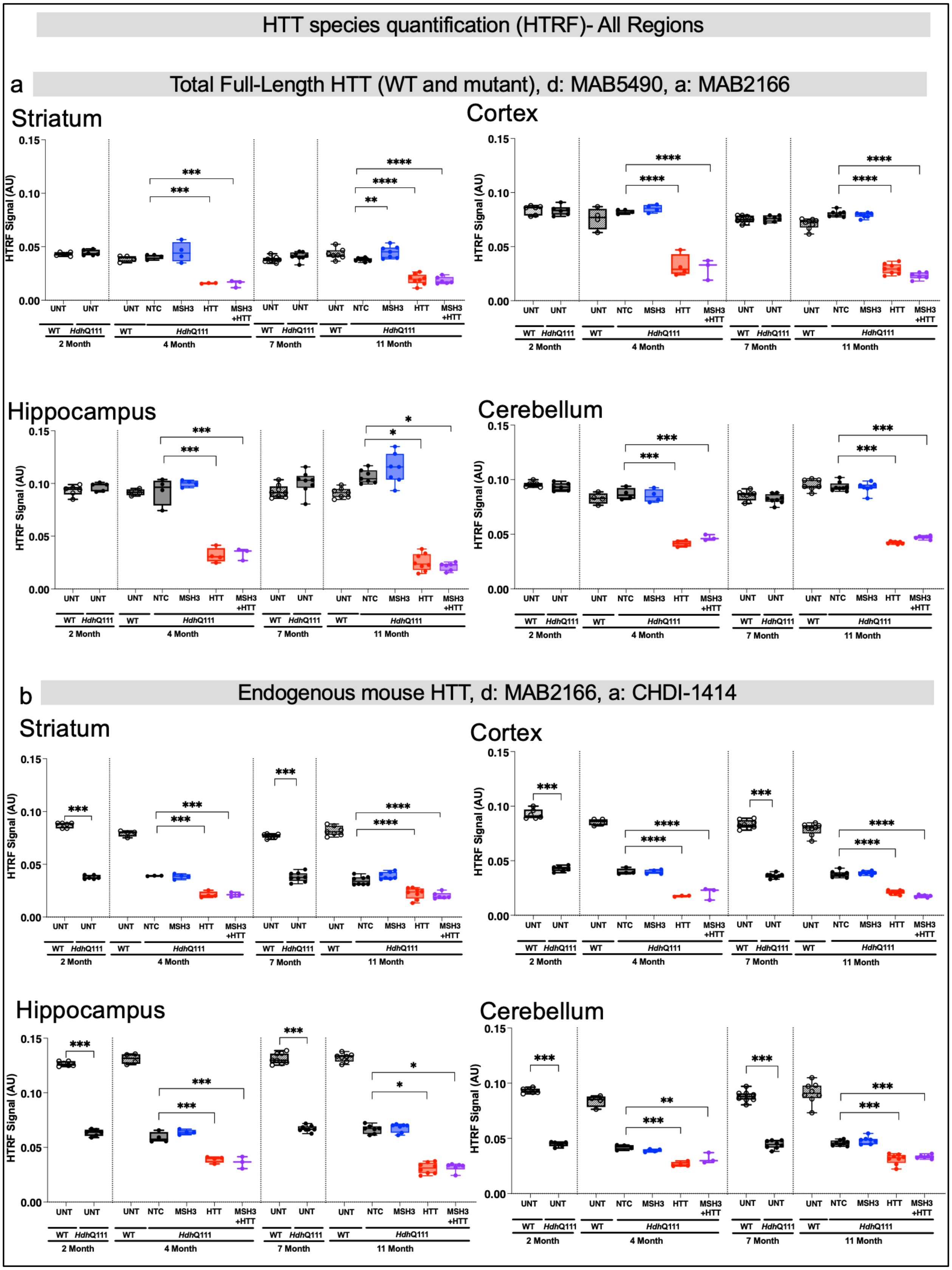
HTT species quantification (HTRF)- All Regions
a
Total Full-Length HTT (WT and mutant), d: MAB5490, a: MAB2166
Striatum
Cortex
Hippocampus
Cerebellum
b
Endogenous mouse HTT, d: MAB2166, a: CHDI-1414
Striatum
Cortex
Hippocampus
Cerebellum
HTRF Signal (AU)
0.15
0.10
0.05
0.00
UNT
NTC
MSH3
HTT
MSH3 +HTT
WT
HdhQ111
2 Month
4 Month
7 Month
11 Month

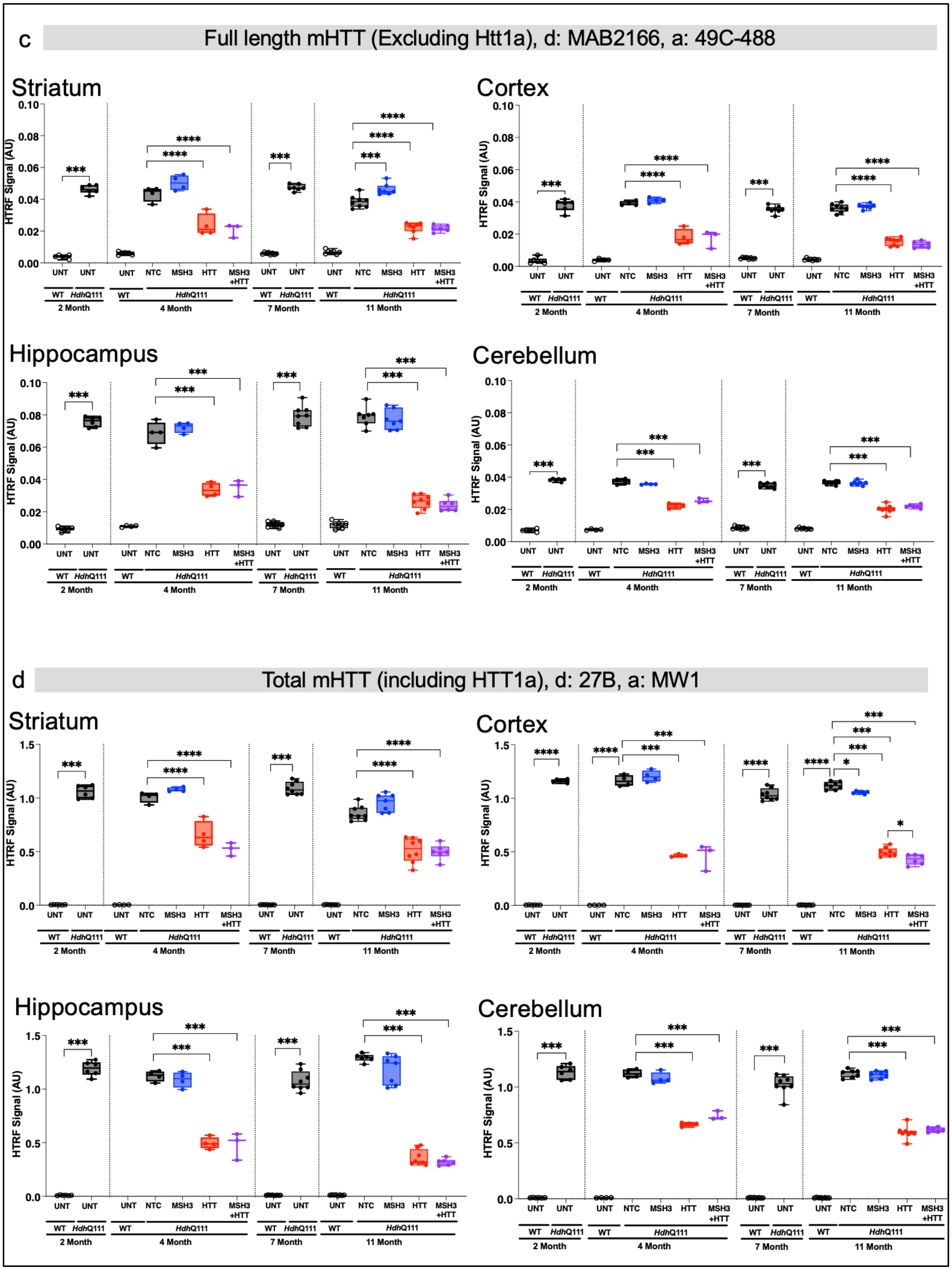
c
Full length mHTT (Excluding Htt1a), d: MAB2166, a: 49C-488
Striatum
Cortex
Hippocampus
Cerebellum
HTRF Signal (AU)
0.00
0.02
0.04
0.06
0.08
0.10
UNT
NTC
MSH3
HTT
MSH3 +HTT
WT
HdhQ111
2 Month
4 Month
7 Month
11 Month
***
****
d
Total mHTT (including HTT1a), d: 27B, a: MW1
Striatum
Cortex
Hippocampus
Cerebellum
HTRF Signal (AU)
0.0
0.5
1.0
1.5
*

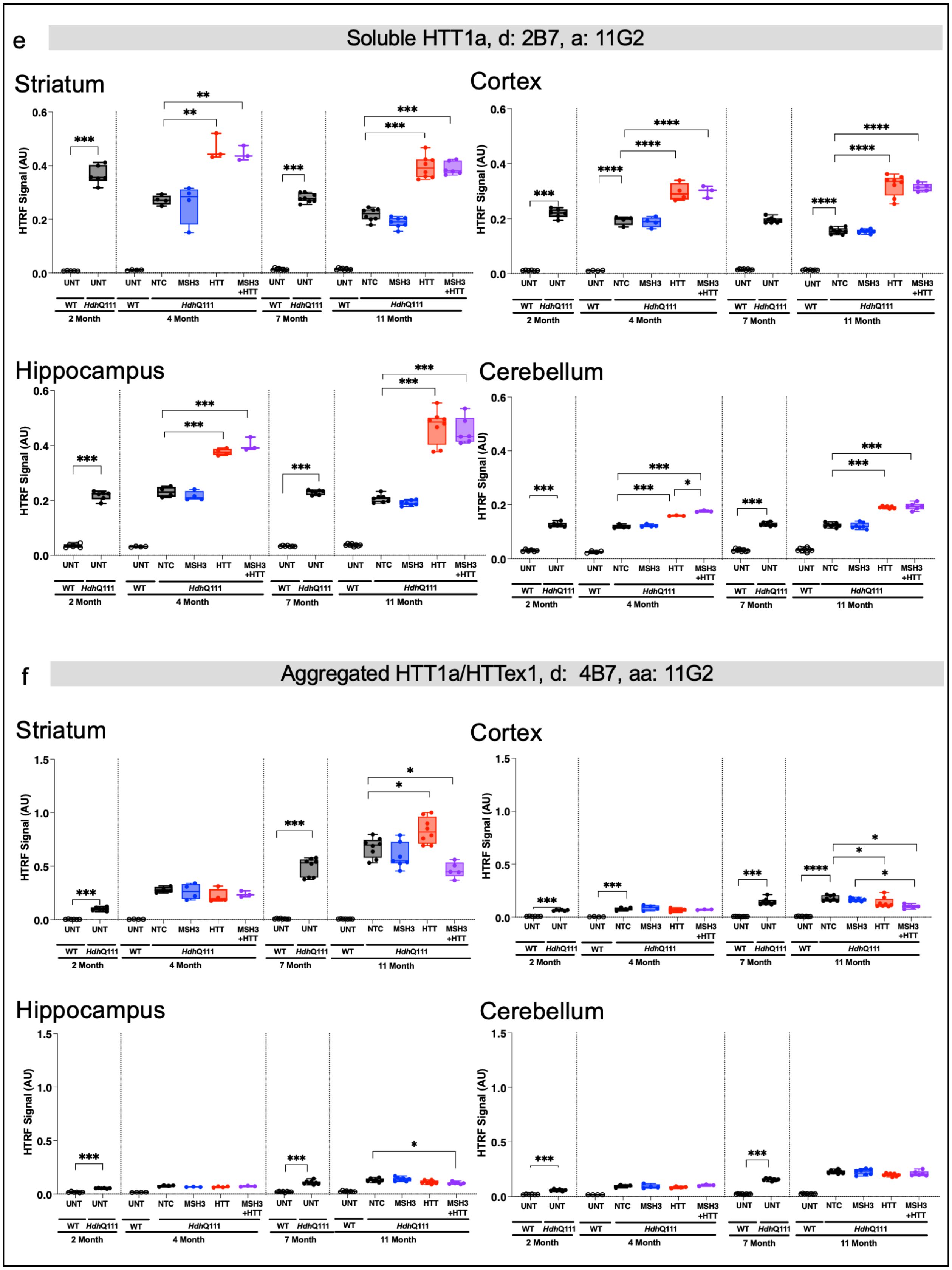

e
Soluble HTT1a, d: 2B7, a: 11G2
Striatum
Cortex
Hippocampus
Cerebellum
HTRF Signal (AU)
UNT
NTC
MSH3
HTT
MSH3 +HTT
WT
HdhQ111
2 Month
4 Month
7 Month
11 Month
f
Aggregated HTT1a/HTTex1, d: 4B7, aa: 11G2
Striatum
Cortex
Hippocampus
Cerebellum
HTRF Signal (AU)

**Supplementary Fig. 5. HTT protein species levels across the striatum, cortex, hippocampus, and cerebellum using homogenous time-resolved fluorescence** (d: donor, a: acceptor). (**a**) Total HTT (d: MAB5490, a: CHDI-2166) (**b**) Endogenous Mouse Htt (d: MAB2166-Tb, a: 1414) (**c)** Soluble mutant HTT (excluding HTT1a) (d: MAB2166-Tb, a: 4C9-488) (**d**) Total soluble mutant HTT (d: 2B7-Tb, a: MW1-d2) (**e**) Soluble HTT1a (d: 2B7-Tb, a: 11G2-d2) (**f**) Aggregated HTT1a (d:4C9-Tb, a:11G2-d2). The X-axis is the genotypes, treatment group, and age of mice at tissue collection. Each dot is data from one mouse. Mice per group at two-month timepoint: WT n=6, Q111 n=6; Mice per group at 4-month timepoint: WT untreated n=4, NTC siRNA n=4, MSH3 siRNA n=4, HTT siRNA n=4, MSH3+HTT siRNA n=3. Mice per group at 7 month timepoint: WT untreated n=8, Q111 untreated n=8, 11-month endpoint: WT untreated n=7, NTC siRNA n=8, MSH3 siRNA n=7, HTT siRNA n=7, MSH3+HTT siRNA n=6. Statistics are one-way ANOVA with all comparisons and Tukey's multiple comparison test. * $p < 0.05$; ** $p < 0.01$; *** $p < 0.001$; **** $p < 0.0001$.

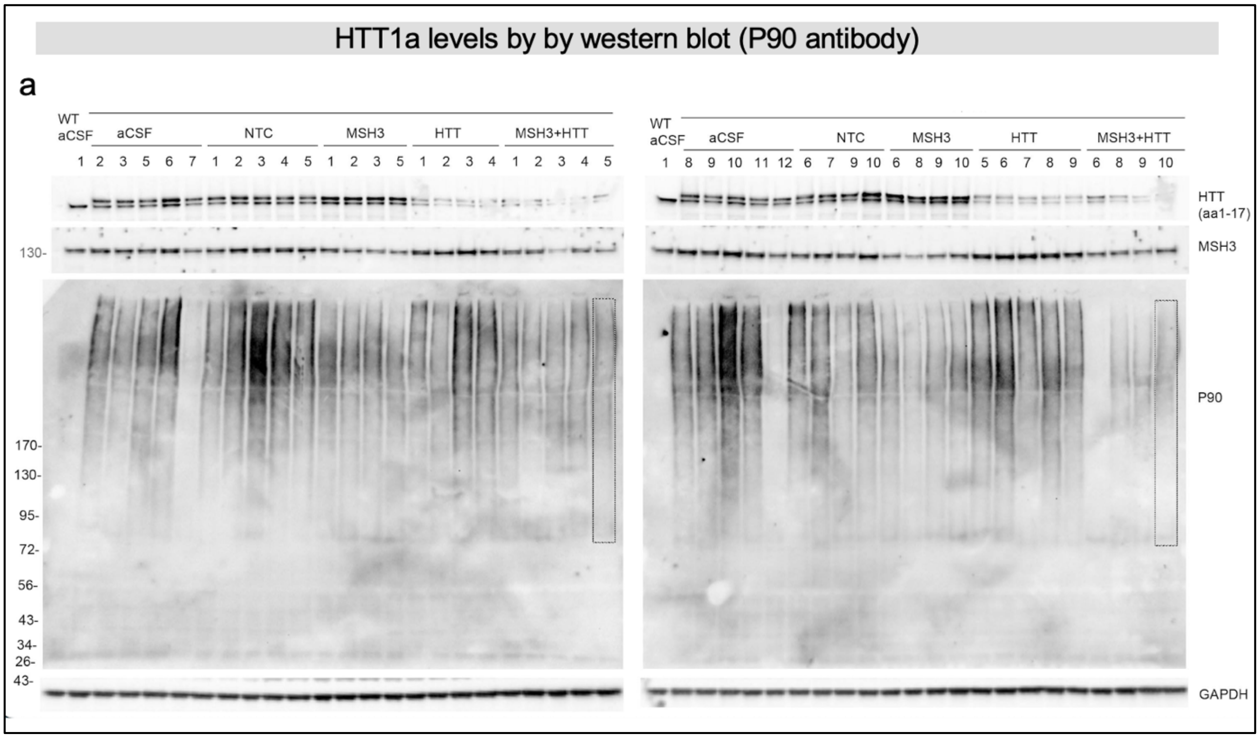


**Supplementary Fig. 6. A combination of blocked somatic expansion and HTT lowering reduces HTT1a protein levels in 12-month-old Q111 mice.** (**a**) Western blot was used to quantify HTT1a levels in Figure 4g.

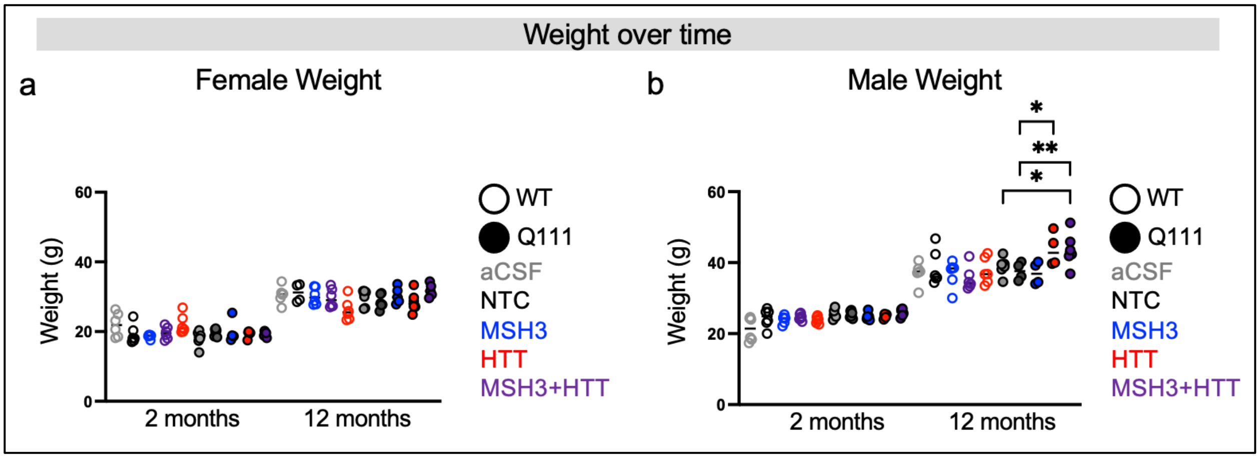


**Supplementary Fig. 7. Q111 mice gain weight similar to WT mice.** Mice weight at age 2 and 12 months. **(a)** Female cohorts **(b)** Male cohorts. Open circle at WT, filled circle are Q111 mice. Age on x-axis, Weight in grams on y-axis. Statistics are one-way ANOVA with Šídák's multiple comparisons test.

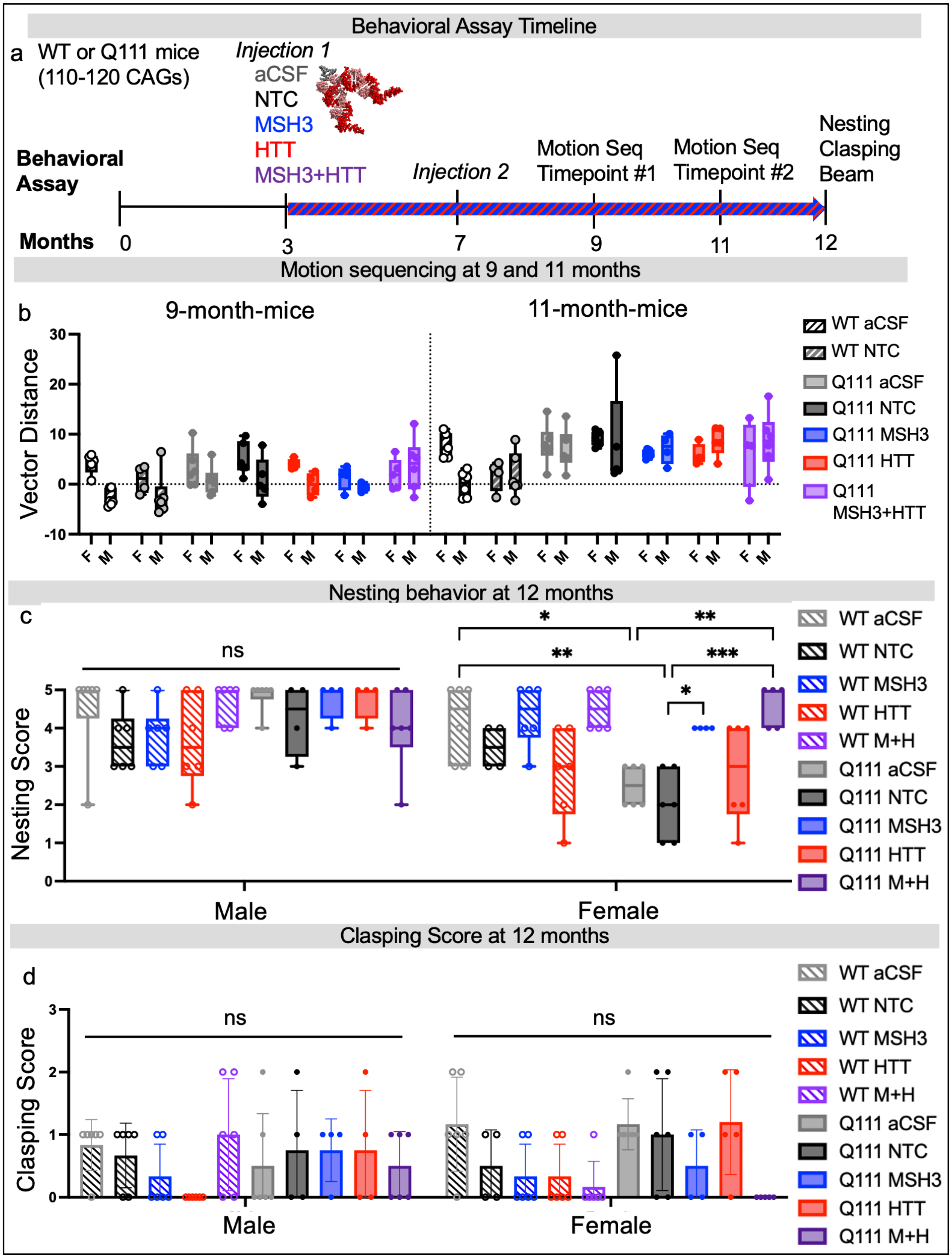

Behavioral Assay Timeline
a
WT or Q111 mice (110-120 CAGs)
Injection 1
aCSF
NTC
MSH3
HTT
MSH3+HTT
Injection 2
Motion Seq Timepoint #1
Motion Seq Timepoint #2
Nesting
Clasping
Beam
Behavioral Assay
Months
0
3
7
9
11
12
Motion sequencing at 9 and 11 months
b
9-month-mice
11-month-mice
Vector Distance
WT aCSF
WT NTC
Q111 aCSF
Q111 NTC
Q111 MSH3
Q111 HTT
Q111 MSH3+HTT
Nesting behavior at 12 months
c
Nesting Score
ns
*
**
**
***
*
Male
Female
WT aCSF
WT NTC
WT MSH3
WT HTT
WT M+H
Q111 aCSF
Q111 NTC
Q111 MSH3
Q111 HTT
Q111 M+H
Clasping Score at 12 months
d
Clasping Score
ns
ns
Male
Female
WT aCSF
WT NTC
WT MSH3
WT HTT
WT M+H
Q111 aCSF
Q111 NTC
Q111 MSH3
Q111 HTT
Q111 M+H

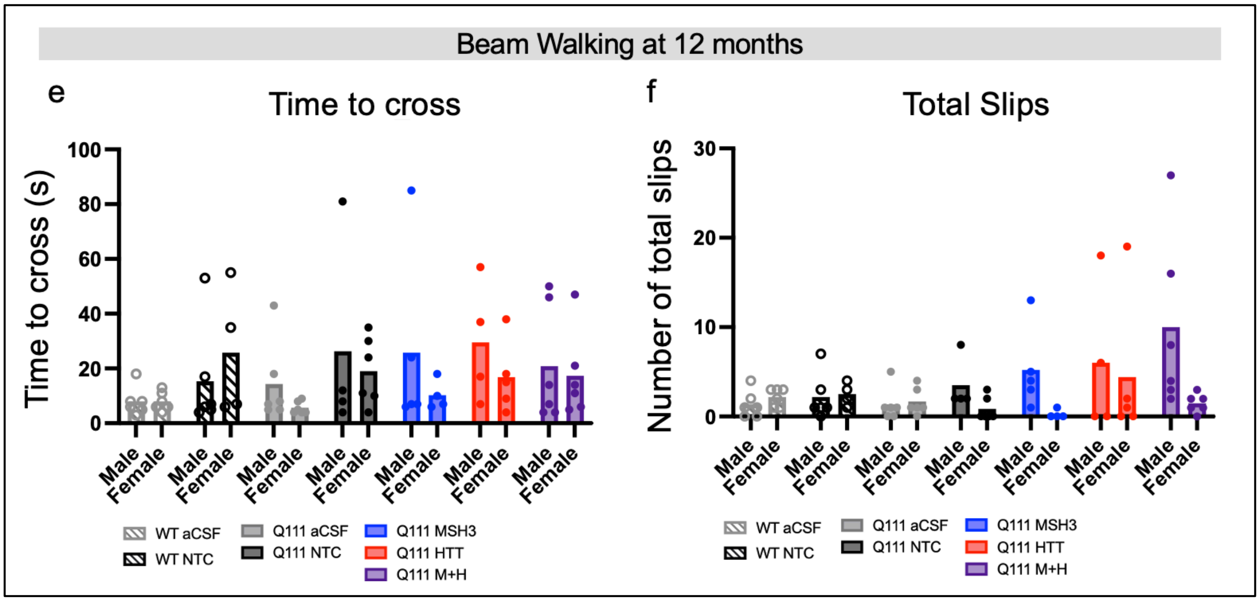


**Supplementary Fig. 8. Female mice show nesting deficits rescued by blocking somatic expansion with MSH3 or MSH3+HTT siRNA treatment.** Motor behavioral assessments in Q111 mice show minimal changes between WT and Q111 mice at 9, 11, or 12 months. (**a**) Experimental design and behavioral analysis performed. (**b**) Data is quantified actions performed in an open field assay[8]. Along top axis, 9 and 11 months refer to mouse age at the time of analysis. WT and Q111 mice were treated with artificial CSF (aCSF) or divalent siRNA programmed with sequences targeting a non-targeting control (NTC), MSH3, HTT, or both MSH3 and HTT. Data represents a correction across genotype vector with 9-month-old aCSF-treated WT and Q111 cohorts used to form a "behavior progression" vector. Treatment cohorts are assessed by where they fall on the defined vector. Statistics are one-way ANOVA with Tukey's multiple comparisons. (**c**) Nesting assay scores in WT and Q111 cohorts at 12 months separated by male and female mice. 5= complete nest formed, 1= nestlet untouched after 24 hours. Statisics are one-way ANOVA with Tukey's multiple comparisons test. (**d**) Clasping assessment in 12-month-old WT or Q111 cohorts separated by male and female mice. Each dot is the performance of one mouse scored over 30 seconds. (**e-f**) WT and Q111 mice across treatment cohorts assessed for beam walking (**e**) time or (**f**) slips. Performance recorded at age 12 months separated by male and female mice. Each dot is the best mouse performance out of three runs. N=9-12 (4-6 males and 4-6 females/group) mice per group for all assays. Performance was assessed blinded to genotype and treatment.

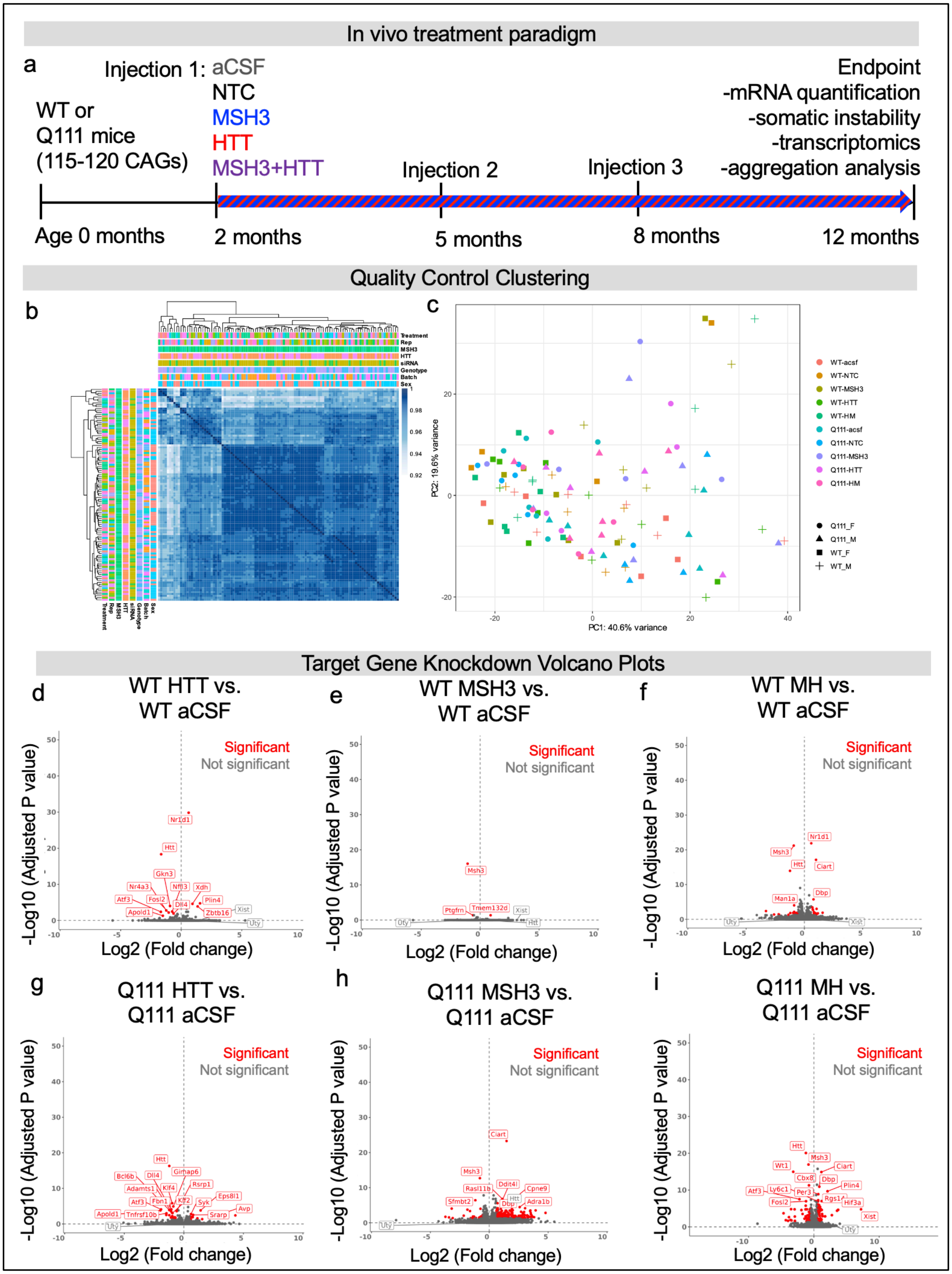
In vivo treatment paradigm
a
Injection 1: aCSF
NTC
MSH3
HTT
MSH3+HTT
WT or
Q111 mice
(115-120 CAGs)
Injection 2
Injection 3
Endpoint
-mRNA quantification
-somatic instability
-transcriptomics
-aggregation analysis
Age 0 months
2 months
5 months
8 months
12 months
Quality Control Clustering
b
c
PC1: 40.6% variance
PC2: 19.6% variance
Target Gene Knockdown Volcano Plots
d
WT HTT vs.
WT aCSF
e
WT MSH3 vs.
WT aCSF
f
WT MH vs.
WT aCSF
g
Q111 HTT vs.
Q111 aCSF
h
Q111 MSH3 vs.
Q111 aCSF
i
Q111 MH vs.
Q111 aCSF
Significant
Not significant
-Log10 (Adjusted P value)
Log2 (Fold change)

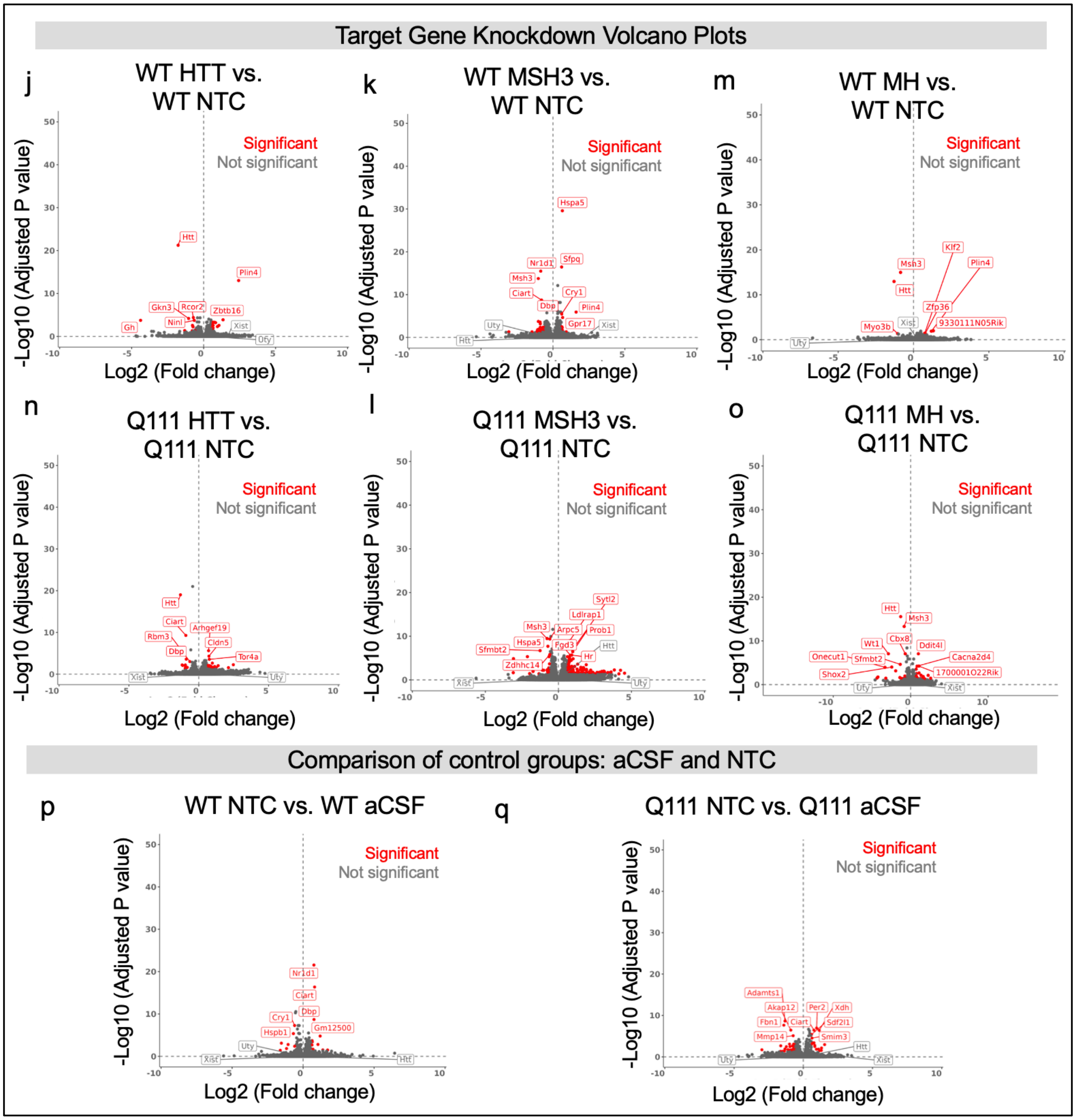


**Supplementary Fig. 9. Bulk striatum transcriptomic data validates on-target knockdown without confounding batch effects.** (**a**) Treatment paradigm of mice (same cohorts at Figures 1-3) used for bulk RNA-sequencing. (**b**) Heat map and (**C**) principal component analysis of all samples across genotypes and treatment groups. (**d-i**) Volcano plot of gene expression changes in the following pairwise comparisons to validate target gene knockdown in aCSF control cohorts:

(**d**) WT HTT versus WT aCSF, (**e)** WT MSH3 versus WT aCSF, (**f**) WT MSH3+HTT versus WT aCSF (**g**) Q111 HTT versus Q111 aCSF, (**h**) Q111 MSH3 versus Q111 aCSF (**i**) Q111 MSH3+HTT versus Q111 aCSF. Red indicates significant gene expression change. (**j-o**) Volcano plot of gene expression changes in the following pairwise comparisons to validate target gene knockdown in NTC control cohorts: (**d**) WT HTT versus WT aCSF, (**e**) WT MSH3 versus WT NTC, (**f**) WT MSH3+HTT versus WT NTC (**g**) Q111 HTT versus Q111 NTC, (**h**) Q111 MSH3 versus Q111 NTC (**i**) Q111 MSH3+HTT versus Q111 NTC. Red indicates significant gene expression change; gray is not significant gene expression change. Significance determined by fold change > 1.5, FDR < 0.05.

Prevention, reversal, recuse (PuRR) classification of siRNA treatment reversal

a

| Full HD Signature | HD Signature | Rev. | Pct. Reversed | Exac. | Pct. Exac. | Full Reversal | Partial Reversal | Neg. Reversal | Super Reversal |
|---|---|---|---|---|---|---|---|---|---|
| HTT vs. Vehicle | 1545 | 191 | 12 | 71 | 5 | 112 | 72 | 1283 | 7 |
| MSH3 vs. Vehicle | 1545 | 372 | 24 | 74 | 5 | 195 | 124 | 1099 | 53 |
| MSH3+HTT vs. Vehicle | 1545 | 565 | 37 | 14 | 1 | 280 | 260 | 966 | 25 |

b

| STR266-R HD Signature | STR-266 Signature | Rev. | Pct. Reversed | Exac. | Pct. Exac. | Full Reversal | Partial Reversal | Neg. Reversal | Super Reversal |
|---|---|---|---|---|---|---|---|---|---|
| HTT vs. Vehicle | 226 | 10 | 4 | 6 | 3 | 4 | 6 | 210 | 0 |
| MSH3 vs. Vehicle | 226 | 48 | 21 | 8 | 4 | 20 | 27 | 170 | 1 |
| MSH3+HTT vs. Vehicle | 226 | 91 | 40 | 0 | 0 | 22 | 68 | 135 | 1 |

**Supplementary Fig. 10. Blocking somatic expansion with HTT lowering using an MSH3+HTT siRNA combination synergistically recused HD signatures in Q111 mice.** Analysis is the prevention, reversal, rescue (PuRR) classification. Differential expression and degree of rescue of genes in the (a) HD signatures or the (b) STR266 signature. Vehicle signature is defined by the overlap of differentially expressed genes in both artificial CSF (aCSF) and non-targeting control (NTC) Q111 cohorts (aCSF-NTC vehicle signature). STR266 list defined by Marchionini et al. 2022. Rev, reversed; Pct, percent; Exac, exacerbation.

| Divalent siRNA sequences used in vivo | | |
|---|---|---|
| **Target** | **Antisense strand** | **Sense strand** |
| **MSH3** | V(mU)#(fG)#(mC)(fA)(fG)(fU)(mU)(fU)(mC)(fA)(mG)(fU)(mU)(fU)#(mG)#(fC)#(mU)#(mU)#(mC)#(fA)#(mU) | (mG)#(mC)#(mA)(fA)(mA)(fC)(mU)(fG)(mA)(fA)(mA)(mC)(mU)(fG)#(mC)#(mA)-DIO |
| **HTT** | V(mU)#(fU)#(mA)(fA)(fU)(fC)(mU)(fC)(mU)(fU)(mU)(fA)(mC)(fU)#(mG)#(fA)#(mU)#(mA)#(mU)#(fA)#(mU) | (mU)#(mC)#(mA)(fG)(mU)(fA)(mA)(fA)(mG)(fA)(mG)(mA)(mU)(fU)#(mA)#(mA)-DIO |
| **NTC** | V(mU)#(fA)#(mA)(fU)(fC)(fG)(mU)(fA)(mU)(fU)(mU)(fG)(mU)(fC)#(mA)#(fA)#(mU)#(mC)#(mA)#(fU)#(mU) | (mU)#(mU)#(mG)(fA)(mC)(fA)(mA)(fA)(mU)(fA)(mC)(mG)(mA)(fU)#(mU)#(mA)-DIO |

**Supplementary Table 1: Divalent siRNA sequences used in vivo**

| Antibody Pairing | Donor (ng / well) | Acceptor (ng / well) | Lysate Dilution Concentration |
|---|---|---|---|
| MAB2166-Tb : CHDI-90001414-d2 | 1 ng | 40 ng | 10% |
| 2B7-Tb : MW1-d2 | 1 ng | 40 ng | 2.5% (1 in 4) |
| MAB2166-Tb : 4C9-488 | 1 ng | 20 ng | 5% (1 in 2) |
| 2B7-Tb : CHDI-90004291-d2 (Clone 11G2) | 1 ng | 20 ng | 10% |
| MAB5490-Tb : MAB2166-d2 | 1 ng | 20 ng | 5% (1 in 2) |
| 4C9-Tb : CHDI90004291-d2 (Clone 11G2) | 1 ng | 30 ng | 10% Crude Lysate |

**Supplemental Table 2: Antibody and lysate concentrations for HTRF assays.** A schematic of these antibodies and their binding location in the HTT protein can be found in Landles et al. 2021
